# Predicting compound activity from phenotypic profiles and chemical structures

**DOI:** 10.1101/2020.12.15.422887

**Authors:** Juan C. Caicedo, Nikita Moshkov, Tim Becker, Kevin Yang, Peter Horvath, Vlado Dancik, Bridget K. Wagner, Paul A. Clemons, Shantanu Singh, Anne E. Carpenter

## Abstract

Recent advances in deep learning enable using chemical structures and phenotypic profiles to accurately predict assay results for compounds virtually, reducing the time and cost of screens in the drug-discovery process. We evaluate the relative strength of three high-throughput data sources—chemical structures, images (Cell Painting), and gene-expression profiles (L1000)—to predict compound activity using a sparse historical collection of 16,186 compounds tested in 314 assays for a total of 679,819 readouts. All three data modalities can predict compound activity with high accuracy in 7-8% of assays tested; replacing million-compound physical screens with computationally prioritized smaller screens throughout the pharmaceutical industry could yield major savings. Furthermore, the three profiling modalities are complementary, and in combination they can predict 18% of assays with high accuracy, and up to 59% if lower accuracy is acceptable for some screening projects. Our study shows that, for many assays, predicting compound activity from phenotypic profiles and chemical structures could accelerate the early stages of the drug-discovery process.

## Introduction

Drug discovery is very expensive and slow. To identify a promising treatment for specific disease conditions, the theoretical landscape of possible chemical structures is prohibitively large to test in physical experiments. Pharmaceutical companies synthesize and test many millions of compounds, yet even these represent a small fraction of possible structures. Furthermore, although complex phenotypic assay systems have proven extremely valuable for identifying useful drugs for diseases where an appropriate protein target is unknown ^1–3^, their reliance on expensive or limited-supply biological materials, such as antibodies or human primary cells, often hinders their scalability.

What if computational models could predict the results of hundreds of expensive assays across millions of compounds at a fraction of the cost? Predictive modeling shows some promise. Most attempts so far have used various representations of chemical structure alone to predict assay activity; this requires no laboratory experiments for the compounds to be predicted (neither to synthesize nor test them), so this is dramatically cheaper than physical screens and enables a huge search space. Deep learning in particular has substantially advanced the state of the art in recent years ^4–16^, and was recently used to discover a novel antibiotic ^17^. As impressive as these capabilities are, chemical structures alone do not seem to contain enough information to predict all assay readouts — their performance may be limited by the lack of experimental information revealing how living organisms respond to these treatments.

Considerable improvements might come from augmenting chemical structure-based features with experimental information associated with each small molecule, ideally information available in inexpensive, scalable assays that could be run on millions of compounds once, then used to predict assay results virtually for hundreds of other individual assays. Most profiling techniques, such as those measuring a subset of the proteome or metabolome, are not scalable to millions of compounds. One exception is transcriptomic profiling by the L1000 assay ^18^, which has shown success for mechanism of action (MOA) prediction ^19^, but is untested for predicting assay outcomes.

Image-based profiling is an even less expensive high-throughput profiling technique ^20^ that has shown great success in compound activity prediction. Cell morphology has been shown to be scalable to annotate compound libraries in real time by determining bioactivity, looking for changes with respect to untreated cells ^21^. In a novel study, Simm et al. ^22^ successfully repurposed images from a compound library screen to train machine learning models to predict unrelated assays; their prospective tests yielded up to 250-fold increased hit rates while also improving structural diversity of the active compounds. More recently, Cell Painting ^23,24^ and machine learning have been used to predict the outcomes of other assays as well ^25,26^.

The complementarity and integration of profiling methodologies and chemical structures to predict compound bioactivity holds promise to improve performance, and has been studied in various ways. The relationships between chemical structures and phenotypic profiles (cell morphology and transcriptional profiles) has been investigated to identify rules that connect structural features with bioactivity patterns in high-dimensional profiling data ^27^. Other studies have looked at combinations of profiles, such as integrating imaging and chemical structures to complete assay readouts in a sparse matrix ^28^ or combining L1000 and Cell Painting for MOA prediction ^19^.

In this work, we aim to evaluate the predictive power of chemical structures, cell morphology profiles, and transcriptional profiles, to determine assay outcomes computationally at large scale. This study does not aim to make predictions in specific assays, which may result in anecdotal success, but rather aims to assess the potential of computational models to support experimental designs in future projects. Our goal is to train machine learning models that predict compound bioactivity taking as input high-dimensional encodings of chemical structures combined with two different types of experimentally-produced phenotypic profiles, imaging (Cell Painting assay) and gene expression (L1000 assay) (Figure 1). Our hypothesis is that data representations of compounds and their experimental effects in cells have complementary strengths to predict assay readouts accurately, and that they can be integrated productively to improve compound prioritization in drug-discovery projects.

**Figure 1.**
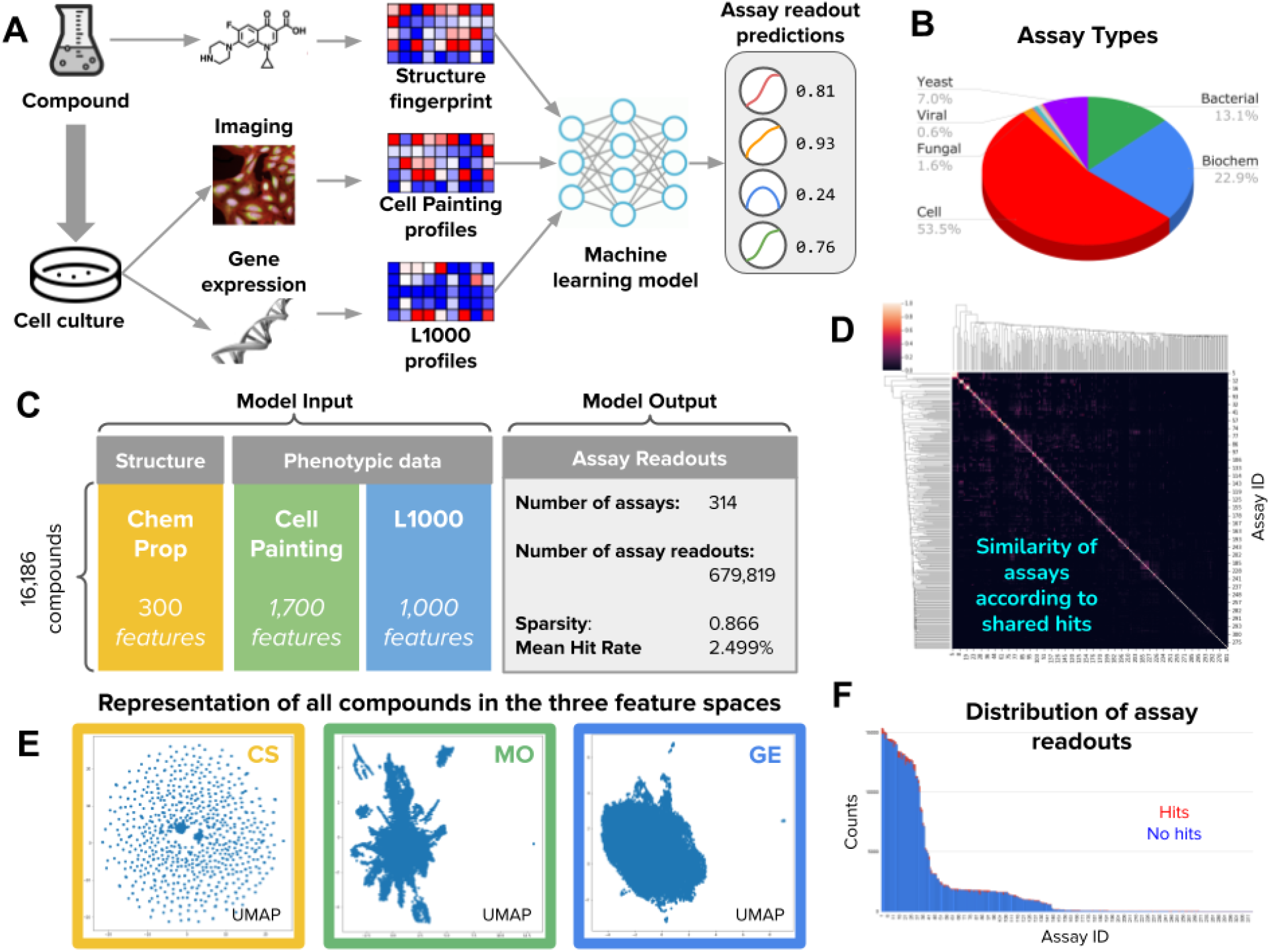
Overview of the workflow and data. A) Workflow of the methodology for predicting diverse assays from perturbation experiments (more details in Supplementary Figures 1 and 2). B) Types of assay readouts targeted for prediction, which include a total of seven categories (Supplementary Figure 14). C) Structure of the input and output data for assay prediction. D) Similarity of assays according to the Jaccard similarity between sets of positive hits. Most assays have independent activity (Supplementary Figure 12). E) UMAP visualizations of all compounds in the three feature spaces evaluated in this study (Supplementary Figure 9). F) Distribution of assay readouts for assays in the horizontal axis sorted by readout counts. The available examples follow a long tail distribution and the ratio of positive hits to negatives (hit rate) is 2.5%

## Results

### Chemical structure, morphology, and gene expression profiles provide complementary information for prediction

We first selected 314 assays performed at the Broad Institute for more than a decade (Figure 1); the assays were not selected based on any metadata and thus are representative of the activity of an academic screening center. Then, we extracted experiment-derived profiles for 16,186 compounds, including gene-expression profiles (GE) from the L1000 assay ^29,30^ and image-based morphological profiles (MO) from the Cell Painting assay ^30,31^. We also computed chemical structure profiles (CS) using graph convolutional nets ^16^ (Figure 1 and Methods). Finally, assay predictors were trained using 5-fold cross-validation using scaffold-based splits (Methods and Supplementary Figures 1 and 9) to quantitatively evaluate the ability of the three data modalities to independently identify hits in the set of held-out compounds.

We found that all three profile types (CS, GE, and MO) can predict a subset of diverse assays with high accuracy, revealing a lack of major overlap among the assays predicted by each profiling modality alone (Figure 2B). This indicates significant complementarity, that is, each profiling modality captures different biologically relevant information. In fact, only three of the 314 assays (<1%) “overlapped” - that is, were accurately predicted using any of the three profiling modalities alone. Chemical structure shares eight well-predicted assays in common with morphology, and none with gene expression, indicating that gene expression captures more independent activity. In fact, gene-expression profiles predicted 14 assays that are not captured by chemical structures or morphology alone, the largest number of *unique* predictors among all modalities (Figure 2B).

**Figure 2.**
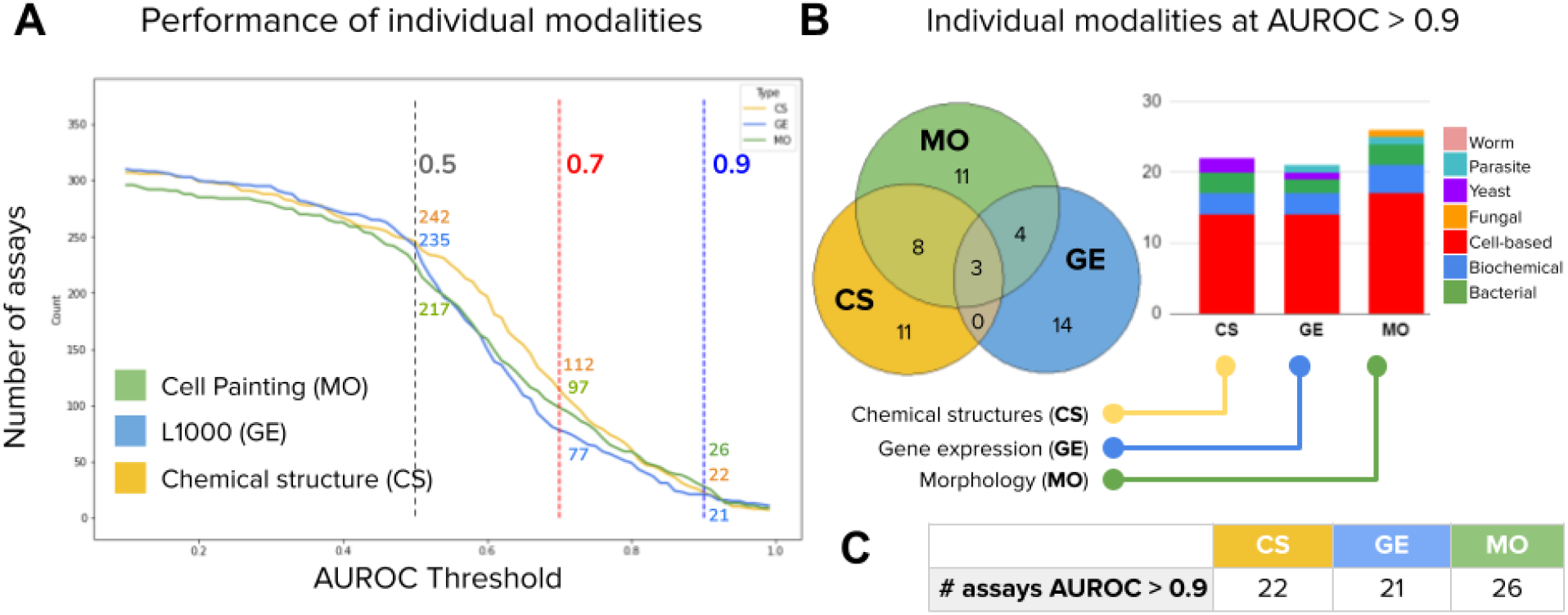
Number of assays that can be accurately predicted using single profiling modalities. All reported numbers are the median result of the five-fold cross-validation experiments run in the dataset. Detailed results of each partition are reported in Supplementary Figure 4 and Supplementary Table 1. A) Performance of individual modalities measured as the number of assays (vertical axis) predicted with AUROC above a certain threshold (horizontal axis). With higher AUROC thresholds, the number of assays that can be predicted decreases for all profiling modalities. We define accurate assays as those with AUROC greater than 0.9 (dashed vertical line in blue). B) The Venn diagrams on the right show the number of accurate assays (median AUROC > 0.9) that are in common or unique to each profiling technique. The bar plot shows the distribution of assay types correctly predicted by single profiling modalities. C) Number of assays well predicted (median AUROC > 0.9) by each individual modality (same as in Figure 3B).

Morphology is able to predict the largest number of assays *individually* (26 vs 22 for CS and 21 for GE) (Figure 2C)., although with AUROC thresholds below 0.8, chemical structure predicts more assays than the other two modalities (Figure 2A). We use the count of predictors with AUROC > 0.9 as our primary evaluation metric, following past studies of assay prediction ^17,19,22^, because it is an indicator of independent bioactivity that is encoded in each profiling modality with a high accuracy threshold. Ideally, profiling modalities would capture sufficient information to accurately predict as many assays as possible; in practice data modalities are limited by the type of bioactivity that they can characterize. The results in Figure 2 reveal the extent to which profiling modalities capture specific bioactivity and confirm that they are indeed mostly different from each other.

### Combining phenotypic profiles with chemical structures improves assay prediction ability

Ideally, combining modalities should leverage their strengths and predict more assays jointly, by productively integrating data. Morphology and gene-expression profiles require wet lab experimentation, whereas chemical structures are always available, even for theoretical compounds, with the only cost being computing their fingerprints. Therefore, we took CS as a baseline and explored the value of adding phenotypic profiles to it.

We first integrated data from different profiling methods using late data fusion and evaluated the performance of combined predictors using the same 5-fold cross validation protocol described for individual profiling modalities. We found that adding morphological profiles to chemical structures yields 27 well-predicted assays (CS+MO) as compared to 22 assays for CS alone (Figure 3C). By contrast, adding gene expression profiles to chemical structures by late data fusion did not increase the number of well-predicted assays as compared to CS alone (21 vs 22 respectively, Figure 3C). This adds evidence that the information provided by GE is more independent from other bioactivity signals. For both phenotypic profiling modalities, early fusion (concatenation of features before prediction) performed worse than late fusion (integration of probabilities after separate predictions, see Methods), yielding fewer predictors with AUROC > 0.9 for all combinations of data types (Supplementary Figure 8 and Supplementary Table 1). The results represent an opportunity for enhancing computational fusion strategies (see Methods - Data fusion).

**Figure 3.**
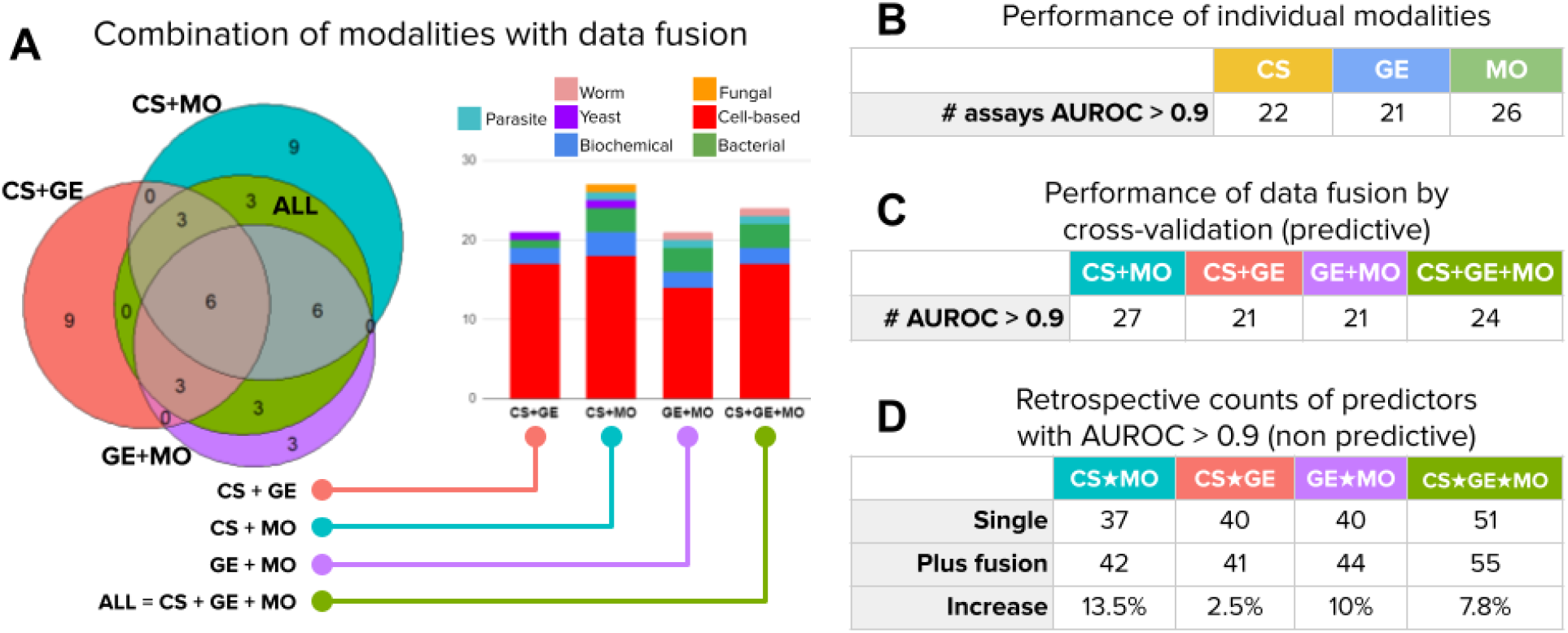
Number of assays that can be accurately predicted using combinations of profiling modalities. Accurate predictors are defined as models with accuracy greater than 0.9 AUROC. We considered all four modality combinations using late data fusion in this analysis: CS+MO (chemical structures and morphology), CS+GE (chemical structures and gene expression), GE+MO (gene expression and morphology), and CS+GE+MO (all three modalities). A) The Venn diagram shows the number of accurately predicted assays that are in common or unique to fused data modalities. The bar plots in the center show the distribution of assay types correctly predicted by the fused models. All counts are the median of results in the holdout set of a five fold cross-validation experiment. B) Performance of individual modalities (same as in Figure 2C). C) The number of accurate assay predictors (AUROC > 0.9) obtained for combinations of modalities (columns) using late data fusion following predictive cross-validation experiments. D) Retrospective performance of predictors. These counts indicate how many assays can be predicted with high accuracy (AUROC > 0.9), whether by single or fused modalities. “Single” is the number of assays reaching AUROC > 0.9 with any one of the specified modalities, i.e., take the best single-modality predictor for an assay in a retrospective way. This count corresponds to the simple union of circles in the Venn diagram in Figure 2B, i.e., no data fusion is involved. “Plus fusion” is the same, except that it displays the number of assays that reach AUROC > 0.9 with any individual or data-fused combination. This count corresponds to the union of circles in the Venn diagram in Figure 2B plus the number of additional assays that reach AUROC > 0.9 when the modalities are fused. For example, the last column counts an assay if its AUROC > 0.9 for any of the following: CS alone, GE alone, MO alone, data-fused CS+GE, data-fused GE+MO, data-fused CS+MO, and data-fused CS+GE+MO. “Increase” indicates the relative increase in the number of assays that can be accurately predicted when using data fusion (calculated from the two rows above it).

Next, we counted the number of unique assays predicted by *any* of the individual profiling modalities using a *retrospective* assessment, which estimates the performance of an ideal data fusion method that perfectly synergizes all modalities. Note that this retrospective assessment is not blind, and simulates a decision maker that chooses the best predictor for an assay after looking at their performance in the hold-out set. It is used here to report the total number of assays that can be successfully predicted using one or another strategy. For example, we found that using the best profiling modality from a given pair can predict around 40 assays (Figure 3D, row “Single”). We use the ★ symbol to denote choosing the best between profiling modalities in retrospect, and the + symbol to denote combining modalities by data fusion.

In retrospect, there are five unique assays that are well predicted using fused CS+MO that could not be captured by either modality alone, indicating complementarity to improve performance for these five assays. Adding them to the list of assays that can be predicted using the single best from CS★MO would yield 42 well-predicted assays total (Figure 3C, row “Plus fusion”), resulting in potential to predict almost twice the assays compared to CS alone (22). Improvements when adding MO to CS were consistently found across other evaluation metrics (AUROC > 0.7 in Supplementary Figure 3 and Supplementary Table 1) and when adding morphological profiles to all other data types and combinations (Figure 3D).

At an AUROC > 0.9, the 42 unique assays that are well predicted with CS★MO represent 13% of the total. An AUROC closer to 0.7 could be acceptable to improve performance and find useful hits in real world projects ^17,22^; we found that for assays with a low baseline hit rate, this accuracy level may be sufficient to increase the ability to identify useful compounds in the screen (Supplementary Figure 3). If a cutoff of AUROC > 0.7 was found to be acceptable, 52% of assays would be well predicted with CS★MO (164 out of 314, Supplementary Figure 3).

The performance of CS★GE also increased the number of assays that CS can predict alone from 22 to 40. There is only one assay that is well predicted using fused CS+GE, which results in 41 unique assays well predicted by both modalities in retrospect. Gene expression also yields similar results when combined with morphology, yielding 40 assays with GE★MO, and predicting four additional assays jointly when using data fusion (GE+MO) for a total of 44 unique assays together.

### Complementarity across all three profiling types

We had hypothesized that data fusion of all three modalities would provide the best assay prediction ability than any individual or subset. However, data-fused CS+GE+MO yielded 24 well-predicted assays (Figure 3C), fewer than could be obtained by data-fused CS+MO (27 assays), which itself was not far from MO alone (26 assays). All of these fall short of the 51 unique assays that, in retrospect, could be identified by taking the single best of any of the three data types CS★MO★GE (Figure 3D). This highlights the need for designing improved strategies for data fusion; early fusion did not improve the situation (Supplementary Figure 8 and Supplementary Table 1).

Likewise, considering the best single, pairwise and all-fused predictors and their combinations, the three data modalities have the potential to accurately predict 55 assays jointly at 0.9 AUROC, not a dramatic improvement compared to 51 unique assays that, in retrospect, could be identified by taking the single best of any of the three data types using CS★MO★GE (Figure 3D). Nevertheless, 55 assays represents 18% of the 314 assays considered in this study. With a threshold of 0.7 AUROC (Supplementary Figure 3), the three modalities can predict 123 assays using data fusion (39% of all 314), and with their retrospective combinations the list grows to 186 assays (59% of all 314). We therefore conclude that if all modalities are available, they are all useful to increase predictive ability.

### Models can predict a diversity of assay types

The morphological and gene-expression profiles used for model training derive from cell-based profiling assays. They can correctly predict compound activity for mammalian cell-based assays, which were the most frequent in this study (Figure 1B, Supplementary Figure 14), but also other assay types, such as bacterial and biochemical (Figure 2B, 3A, Supplementary Figure 14). Still, cell-based assays were the best-predicted by the phenotypic profiles as well as by chemical structures: from 168 cell-based assays, 14, 14 and 17 are accurately predicted by CS, GE and MO respectively (8%, 8%, 10%); by contrast, from 72 biochemical assays, 3, 3 and 4 were predicted by CS, GE and MO respectively (4%, 4%, 5%).

We nevertheless conclude that well predicted assays include diverse assay types, i.e., phenotypic profiling strategies are not constrained to predict cell-based assays only, even though both profiling methods are cell-based assays themselves. Each modality predicted assays in 4-5 of the 9 assay categories when used alone (Figure 2B). When MO is combined with CS, they can predict assays in 6 out of the 9 assay categories included in this study (Figure 3A).

As noted above, only a few assays benefit from combining information of various profiling modalities. We examined four assays with increased fused accuracy more closely (Figure 4). The *Ras selective lethality* assay, a cell-based assay, can be predicted with a maximum AUROC of 0.76 using MO alone, but when the three modalities are combined with data fusion, the performance increases to AUROC 0.88. Similarly, the *Beta Cell Apoptosis Screen assay* reaches an accuracy of 0.88 using MO alone, but when CS is added, performance increases to 0.93 AUROC. These examples indicate that fusing information from various modalities can improve predictive performance, but the fusion result may depend on several factors such as the diversity and availability of training examples and the biology measured by the specific assay.

**Figure 4.**
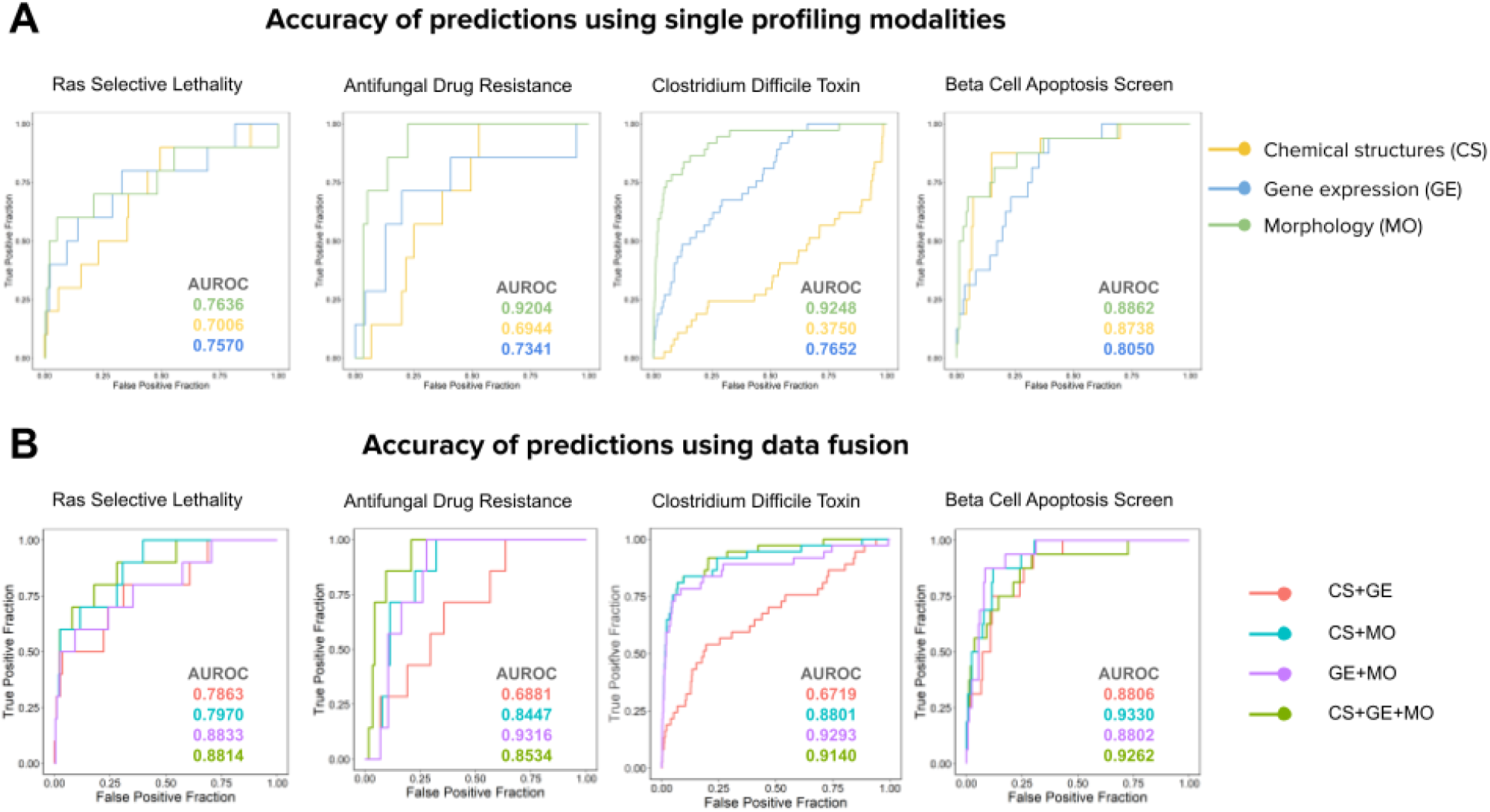
Prediction accuracy of assays where prediction accuracy benefits from fusion. The plots are Receiver Operating Characteristic (ROC) curves and the area under the curve (AUROC) is reported for each curve with the corresponding color. Four assays were selected for panels A and B, in order from left to right: Ras Selective Lethality (cell-based), Antifungal Drug Resistance (fungal), Clostridium Difficile Toxin (bacterial), Beta Cell Apoptosis Screen (cell-based). A) Performance of predictors when using a single profiling method. B) Performance of predictors when using combinations of profiling methods.

### Assay predictors trained with phenotypic profiles can improve hit rates

Predictive modeling using machine learning to reuse phenotypic profiles in a large library of compounds can enable virtual screening to identify candidate hits without physically running the assays. Here, we compare the empirical hit rate of testing a large subset of candidate compounds physically in the lab, vs the hit rate of testing only the top predicted candidates obtained with a computational model (Supplementary Figure 6). The ratio between these two hit rates is what we term *folds of improvement*, a factor indicating the expected experimental efficiency if the computational model identifies relevant compounds to follow up with.

We found that predictors meeting AUROC > 0.9 in our experiments produce on average a 50 to 70-fold improvement in hit rate (i.e., compounds with the desired activity, see Supplementary Figure 7) for assays with a baseline hit rate below 1%. A baseline hit rate below 1% means that hits are rare for such assays, i.e., in order to find a hit we need to test at least 100 compounds randomly selected from the library. Assays with low hit rates are common in real world screens, and therefore, more expensive to run in practice. With computational predictions improving hit rates by 50 fold, the speed and return of investment could be potentially very high. We also note that for assays with higher baseline hit rates (e.g. 10% to 50%), the machine learning models can reach the theoretical maximum fold of improvement by accurately predicting all the hits in the top of the list (Supplementary Figure 7). We conclude that when assay predictors are accurate enough, they can significantly accelerate compound screening and reduce the resources required to identify useful hits.

## Discussion

Predicting bioactivity of compounds could become a powerful strategy for drug discovery in light of ever-improving computational methods (particularly, deep learning) and ever-increasing rich data sources (particularly, from profiling assays). Here, we used the Chemprop software for learning predictors from chemical structures, and to combine the molecular fingerprint with phenotypic profiles obtained from images (Cell Painting) and gene expression (L1000). We conducted this study using baseline feature representations, and arguably, the results could be improved in future research by using alternative chemical structure embeddings ^32–34^, learned image features ^25,35^, or latent spaces for gene expression ^36^.

We discovered that all three profile modalities—chemical structure, morphology, and gene expression—offer independently useful information about perturbed cell states that enables predicting different assays. Chemical structure is always readily available for a given compound. The two profiling modalities that require physical experimentation bring different strengths to the assay prediction problem, and if available, they can be leveraged to run virtual screens to prioritize compound candidates in drug-discovery projects.

In retrospect, we found that data fusion strategies increased the number of well-predicted assays by only 3-10%, depending on the subset of modalities tested, as compared to simply using each profiling modality independently for prediction. We believe this argues for further research on how best to integrate disparate profiling modalities, capturing the strengths of each individually as well as the complementarity of their combinations. Nevertheless, using late data fusion to combine each subset of available modalities does offer some improvement versus each individually and is likely worthwhile given its ease of implementation.

Given the low cost of carrying out Cell Painting, it is practical in many settings to profile an entire institution’s compound library. Then, a modest-sized library of a few thousand compounds would be tested in each new assay of interest, providing sufficient data to assess whether an accurate predictor could be trained on these data, using CS alone, MO alone, or a data-fused combination of CS+MO. Taking into account the baseline hit rate for the assay, researchers could decide whether the predictor will increase the hit rate sufficiently to warrant a virtual screen against a large compound library for which morphological profiles are already available (within an institution, or publicly available profiles ^37^), followed by cherry picking a small set of predicted hits and testing them for actual activity in the assay.

Although we suggest a few thousand compounds for the training set based on the data shown in Supplementary Figure 5, it remains to be fully evaluated how many training points are needed to achieve strong predictivity — in fact, it is likely that the number and structural diversity of hits in the training set more strongly influences predictivity than the total number of assay data points. Nevertheless, in most academic and industry screening centers, preparing a training/test set of ~17,000 compounds, as we used here, is practical. It also remains to be determined what else needs to be evaluated to help decide if an assay is likely to be predictable, which may require additional knowledge of the target and assay type, and characterizing correlations between the bioactivity of interest and profiling modalities, as well as the assay activity distribution.

Based on our results, and depending on whether an AUROC of 0.9 or 0.7 is the threshold for accuracy needed given the baseline hit rate of the assay, 18-59% of assays should be predictable using a combination of chemical structures, morphology and gene expression, saving the time and expense of screening these assays against a full compound library. Especially considering potential improvements in data integration techniques and deep learning for feature extraction, this strategy will be fruitful to accelerate the discovery of useful chemical matter.

## Methods

### Profiling datasets

For this study, we used a compound library of over 30,000 compounds screened in high-throughput ^30^. Of these compounds, about 10,000 came from the Molecular Libraries Small Molecules Repository, another 2,200 were drugs and small molecules, and the remaining 18,000 were novel compounds derived from diversity oriented synthesis. U2OS cells were plated in 384-well plates and treated with these compounds in 5 replicates, using DMSO as a negative control. The Cell Painting and L1000 platforms were used to generate morphological and transcriptional profiling data, respectively, as previously described ^30^.

### Assay readouts

On the assay side, we collected a list of 529 assays from drug discovery projects conducted at the Broad Institute at different scales, and we kept those where at least a subset of the small molecules in the compound library described above was tested. We kept assays that had a non-empty subset of hits identified for training and evaluation. We prepared assay performance profiles following a double sigmoid normalization procedure to ensure that all readouts are scaled in the same range ^38^. Then, we computed the Jaccard similarity of hits between pairs of assays to estimate the common set of compounds detected by them, and then removed assays that measure redundant compound activity (Supplementary Figure 12). That resulted in a final list of 314 assays with their corresponding readout results (Supplementary Figure 11), and the compound-assay matrix had 13.4% of known entries (86.6% sparsity).

### Training / Test splits

The total number of compounds in the library that had the three types of information required to conduct the experiments in our project (Cell Painting images, L1000 profiles, and assay readouts) was 16,979. From this set of compounds we removed frequent hitters, defined as compounds that are positive hits in at least 10% of the assays (hits in 31 assays or more), which removed 24 compounds from the list. We also applied all pan-assay interference (PAINS) filters ^39^ implemented in RDKit, which removed 786 compounds, resulting in 16,186 compounds in the final dataset.

We aimed to evaluate the ability of each data modality to predict assays for chemical structures that are *distinct* relative to training data. This is because there is little practical value to screen for additional, similar structures (scaffolds) to compounds already known to have activity; in drug discovery, any compounds with positive activity undergo medicinal chemistry where small variations in structure are synthesized and tested to optimize the molecule. We therefore report results using cross-validation partitions that ensure that similar classes of structures are not included in both the training and hold-out sets, given that this scheme corresponds to the most practical, real world scenario (Supplementary Figure 9).

We used 5-fold cross-validation using Bemis-Murcko clustering ^40,41^, and assigned clusters to training or test in each fold accordingly. The main experimental design for the results reported in the main text is illustrated in the Supplementary Figure 1. The distribution of chemical structure similarity according to the Tanimoto coefficient metric on Morgan fingerprints (radius=2) is reported in Supplementary Table 10 for each of the 5 cross-validation groups. As additional control tests, we run 5-fold cross-validation experiments following the same design as above but splitting the data according to k-means clusters in the morphology feature space and in the gene-expression space (Supplementary Figure 9 and Supplementary Table 2), as well as a control experiment with fully random splits (Supplementary Table 2).

The control splits based on randomized data as well as the MO and GE modalities were used to check for and identify potential biases in the data. These splits do not have practical applications in the lab, and were used as computational simulations to test the alternative hypothesis that predictors have a disadvantage when the training data are drawn from a distribution that follows similarities in CS, MO or GE. The results in the Supplementary Table 2 indicate that there is no major change in performance when using CS, GE or random splits; however, MO splits reduced performance significantly for all data modalities. This process revealed the need to correct for batch effects in MO data to minimize the influence of technical artifacts. All results presented in the main text were obtained from MO data that has been batch corrected (see Image-based morphological profiles below).

### Representation of chemical structures (CS) using Chemprop

We used the Chemprop software (http://chemprop.csail.mit.edu/) to train directed-message passing neural networks for learning chemical structure embeddings. The software reconstructs a molecular graph of chemicals from their SMILES string representation, where atoms are nodes and bonds are edges. From this graph, a model applies a series of message passing steps to aggregate information from neighboring atoms and bonds to create a better representation of the molecule. For more details about the model and the software, we refer the reader to prior work ^16,17,42^. In addition to learning representations for chemical structures, we used Chemprop to run all the machine learning models evaluated in this work to base all the experiments on the same computational framework. Also, we evaluated the predictive models for CS using learned features as well as Morgan fingerprints computed with the RDKit software (radius=2), and we found that both yield comparable results in our main experiments (Supplementary Table 2, columns CS-GC [Graph Convolutions] and CS-MF [Morgan Fingerprints]).

The representation of chemical structures is learned from the set of ~13,000 training examples, unlike morphological or gene-expression features, which were obtained without learning methods (hand-engineered features). The scaffold split used in our experiments may pose an apparent disadvantage to the learning of chemical structure representations because it may not learn to represent important chemical features in new scaffolds. Previous research by Yang et al. ^16^ has shown that Chemprop can generalize to new scaffolds accurately. In addition, the chemicals may also generate new phenotypes in the morphological and gene-expression space, which are not seen by the models during training, resulting in a fair comparison of representation power among all modalities. We tested the effect of creating partitions with other modalities other than scaffolds from chemical structures, and we discuss these results in the Train / Test splits subsection above as well as in Supplementary Table 2 and Supplementary Figure 9.

### Image-based morphological (MO) profiles from the Cell Painting assay

The Cell Painting assay ^23,24,27,30^ captures fluorescence images of cells using six dyes to label eight major cell compartments. The five-channel, high-resolution images are processed using the CellProfiler software (https://cellprofiler.org/) to segment the cells and compute a set of 1,700+ morphological features at the single-cell level. These features are aggregated into well- and treatment-level profiles that capture the central statistics of the response of cells to the treatment.

Before computing treatment-level profiles, we used the Typical Variation Normalization (TVN) ^43^ transform to correct for batch effects using well-level profiles (see Supplementary Figure 9). TVN is calculated using DMSO control wells from all plates to compute a sphering transform that reduces the data to a white noise distribution by inverting all the non-zero eigenvalues of the matrix. This transformation is later used to project all treatment wells in a new space, where controls have a neutral representation and treatments may have phenotypic variations highlighted. This transform minimizes batch effects by obtaining a feature space where the technical variations sampled from controls are neutralized to enhance the biological signal.

After applying the TVN transform at the well-level profiles, we aggregate them into treatment-level profiles to conduct our assay prediction experiments. Supplementary Figure 8 shows UMAP plots of the morphology data before and after the TVN transformation. In our study, we used treatment-level profiles in all experiments. For more details about Cell Painting ^24^, CellProfiler ^44^, and the profiling steps ^45^, see the corresponding references.

### Gene-expression (GE) profiles from the L1000 assay

The L1000 assay measures transcriptional activity of perturbed populations of cells at high-throughput. These profiles contain mRNA levels for 978 landmark genes that capture approximately 80% of the transcriptional variance ^18^. The assay was used to measure gene expression in U2OS cells treated with the set of compounds in our library. Both the profiles and the tools to process this information are available at https://clue.io/.

### Predictive model

#### Model architecture

The predictive model is a feedforward, fully connected neural network with three layers and ReLU activation functions. This simple architecture takes as input compound features (or phenotypic profiles) and produces as output the hit probabilities for all assays (see Supplementary Figure 8). When the representation of chemical structures is learned, additional layers are created before the predictive model to compute the message passing graph convolutions. These extra layers and their computation follow the default configuration of Chemprop models ^16^ and are only used for chemical structures.

#### Loss and training

The model architecture described above is trained in a multi-task manner ^5^, allocating a binary output for each assay. We used the logistic regression loss function on each assay output and the total loss is the sum over all assays. During training, the model computes this loss for each assay output independently using the available readouts. If the assay readout is not available for some compounds in the mini-batch, these outputs are ignored and not taken into account to calculate gradients. This setup facilitates learning predictive models with sparse assay readouts. We use a mini-batch size of 50 compounds with a sparse matrix of 314 labels, and no explicit class balancing was applied during training.

#### Hyperparameter optimization

The hyperparameters of the network are optimized on the training data for each feature grouping and for each cross-validation round. These parameters are: number of fully connected layers (choice between 1, 2 or 3), dropout rate for all layers (between 0 and 1), and hidden layer dimensionality (if applicable, between 100 and 2,500). The best parameters are identified by further splitting the training set in three parts, with proportions 80% for training, 10% for validation and 10% for reporting hyperparameter optimization performance. Then, these parameters are used to train a final model that is used to make predictions in the hold-out partition of the corresponding cross-validation set.

### Data fusion

The input to the neural network can be the features of one or all modalities used in our experiments. To combine features from multiple data modalities, we used two strategies (Supplementary Figure 8): A) early data fusion, where feature vectors from two or three modalities are concatenated into a single vector. B) Late data fusion, where each modality is used to train a separate model, and then the prediction scores for a new sample are aggregated using the maximum operator. Our results show that, despite its simplicity, late data fusion works best in practice (see Supplementary Table 2), but the results also suggest that more research needs to be done to effectively combine multiple data modalities.

Combining disparate data modalities (sometimes called multimodal or multi-omic data integration) is an unmet computational challenge especially when not all the assays can be accurately predicted. Our results indicate that the three data modalities only predict a small fraction of the assays in common (Figure 2B, only three assays are predicted by all modalities), suggesting that in most cases, at least one of the data modalities will effectively introduce noise for predicting a given assay. When one of the data modalities cannot signal the bioactivity of interest, the noise-to-signal ratio in the feature space increases, making it more challenging for predictive models to succeed. This explains why late fusion, which independently looks at each modality, tends to produce better performance.

### Performance metrics

To evaluate the performance of assay predictors we used the area under the receiving operating characteristic (ROC) curve, also known as the AUROC metric, which has a baseline random performance of 0.5. During the test phase, we run the model over all compounds in the test set to obtain their hit probabilities for all assays. With these probabilities, we compute AUROC for each assay using only the compounds that have ground truth annotations (either positive hits or negative results), and we ignore the rest of the compounds that have no annotation for that assay (unknown result or compound never tested).

We define a threshold of AUROC > 0.9 to identify assays that can be accurately predicted, and with this threshold, our second performance metric is focused on counting how many assays, from the list of 314 in our study, can be accurately predicted. For comparison, we also calculated Average Precision (AP) and area under the precision-recall curve (AUPRC) which are reported in Supplementary Tables 1, 2 and 3.

In addition, we measured hit-rate improvement for individual assays as the ratio between the hit rate obtained using the computational predictors and the hit rate observed in the lab (the “baseline” hit rate):

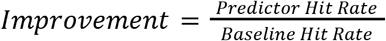

*Predictor hit rates* are calculated as the proportion of positive hits observed in the top 1% of the ranked list of predictions, while *baseline hit rates* are calculated as the number of hits identified in the complete set of compounds tested for that assay in the original experiment. For an illustration of this performance metric see Supplementary Figure 6 and Supplementary Figure 7 for the results.

## Data and code availability

The morphological and gene-expression profiles were originally created and published by Wawer, M. J. et al. ^30^, and can be downloaded from: http://www.broadinstitute.org/mlpcn/data/Broad.PNAS2014.ProfilingData.zip

The Cell Painting images were also made available by Bray et al. ^31^, and can be obtained from the following link: http://gigadb.org/dataset/100351

The latest version of morphological profiles is also available in the following AWS S3 bucket: https://registry.opendata.aws/cell-painting-image-collection/

The Chemprop software and source code used for training machine learning models can be found in the following link: http://chemprop.csail.mit.edu/

The analysis code to reproduce the experiments reported in the paper can be found in the following link: https://github.com/carpenterlab/puma_project

Anonymized assay data to reproduce the analysis in the paper will be made available soon.

## Acknowledgements

This study was supported by a grant from the National Institutes of Health (R35 GM122547 to AEC), and by the Broad Institute Schmidt Fellowship program (JCC). The authors are grateful for guidance and thoughtful discussions with Regina Barzilay and Tommi Jaakkola which improved the analysis and manuscript.

## Supplementary Material

### Experimental design

**Supplementary Figure 1.**
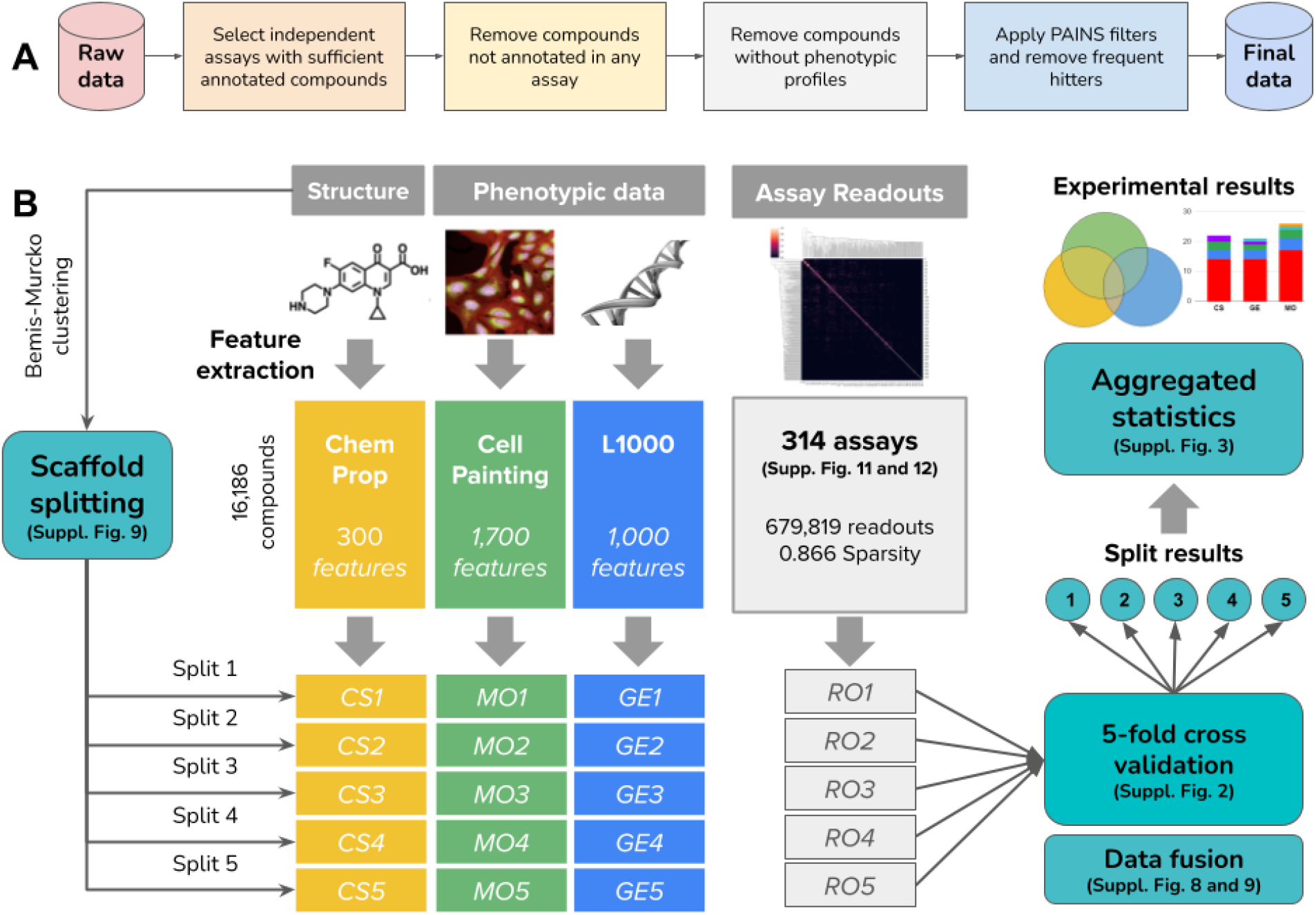
Illustration of the experimental design in this study. A) Data selection and filtering pipeline to construct the dataset used in this study. The process is linear and the order of steps is followed one at a time. We first select 314 assays from more than 500 available (see Supplementary Figure 11 and 12), and with those targets fixed, we proceed to clean the list of compounds with various other filters. B) We considered the problem of assay prediction from three compound representations: features of the chemical structure, and phenotypic features of the effect of compounds measured by imaging (Cell Painting) and gene expression (L1000). We conducted a 5-fold cross-validation experiment splitting the compounds in 5 groups according to scaffold similarity using the Bemis-Murcko clustering. The profiles for compounds in each of these groups were separated together with the corresponding assay readouts. The training of models and test of predictions is carried out independently for each fold, and the results are aggregated to generate summarized statistics of the experimental results.

**Supplementary Figure 2.**
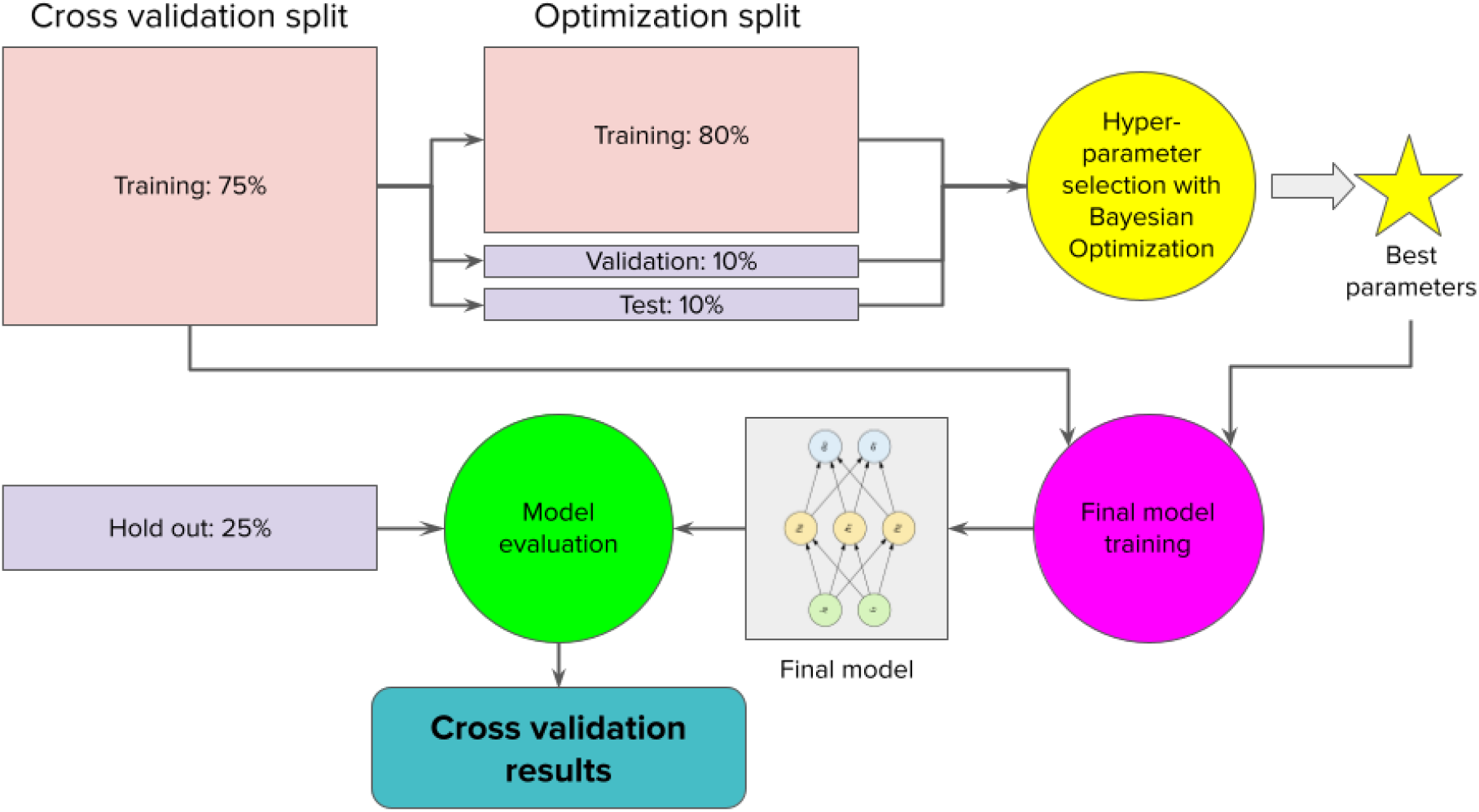
Pipeline of cross-validation experiments. The models trained and evaluated in our experiments are conducted following this protocol: for each split in the 5-fold validation scheme, we take the training dataset and split it again in three parts: 80% for training, 10% for validation and 10% for testing. In this partition, we run hyperparameter search using Bayesian optimization to calibrate the parameters described in the Methods section, subsection Predictive model and data fusion. The Bayesian optimization model uses the 10% assigned for validation to search better parameters at each iteration, and when the search is complete, a final evaluation is performed on the 10% test set with a subset of the best candidates to identify the hyperparameters with better out of sample generalization. These best hyperparameters are used to train a final model with the entire training data in the original split, which is later evaluated with the subset held out for test. The results out of this evaluation are reported in the main text as well as in the rest of the manuscript.

### Additional results

**Supplementary Figure 3.**
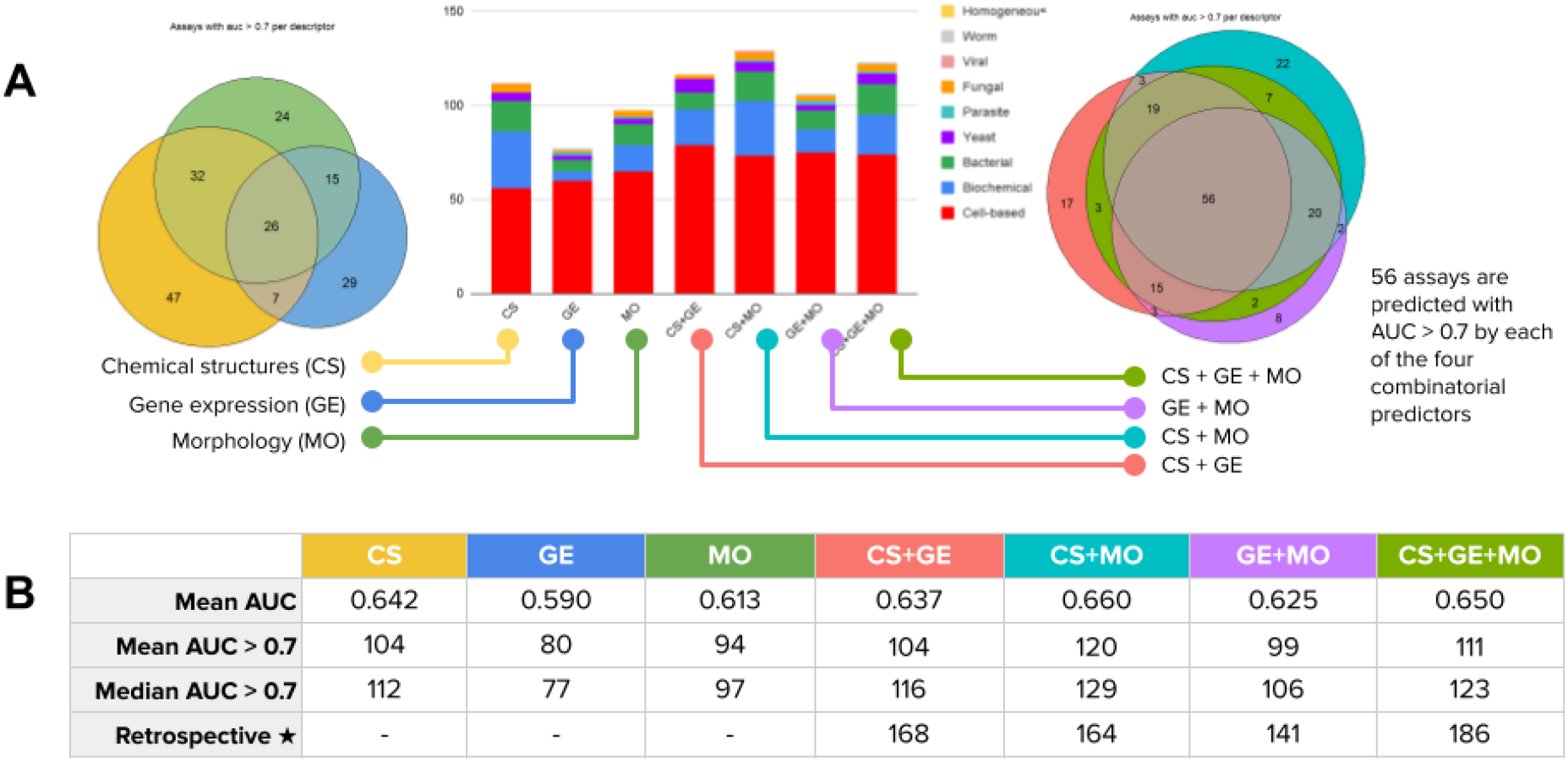
Summary of the number of assays predicted with models that have AUROC > 0.7, which is a lower performance threshold than the one used throughout our study. The total number of acceptable assay predictors increases when the threshold is lower, and chemical structures can yield more predictors that meet this level of performance. Importantly, predictors that reach performance above 0.7 AUROC are also capable of improving hit rates in many cases (see yellow points in Supplementary Figure 7). The row “Retrospective” in Table B presents the number of assays with AUROC > 0.7 that would be predicted by any of the modalities individually or their combinations.

**Supplementary Figure 4.**
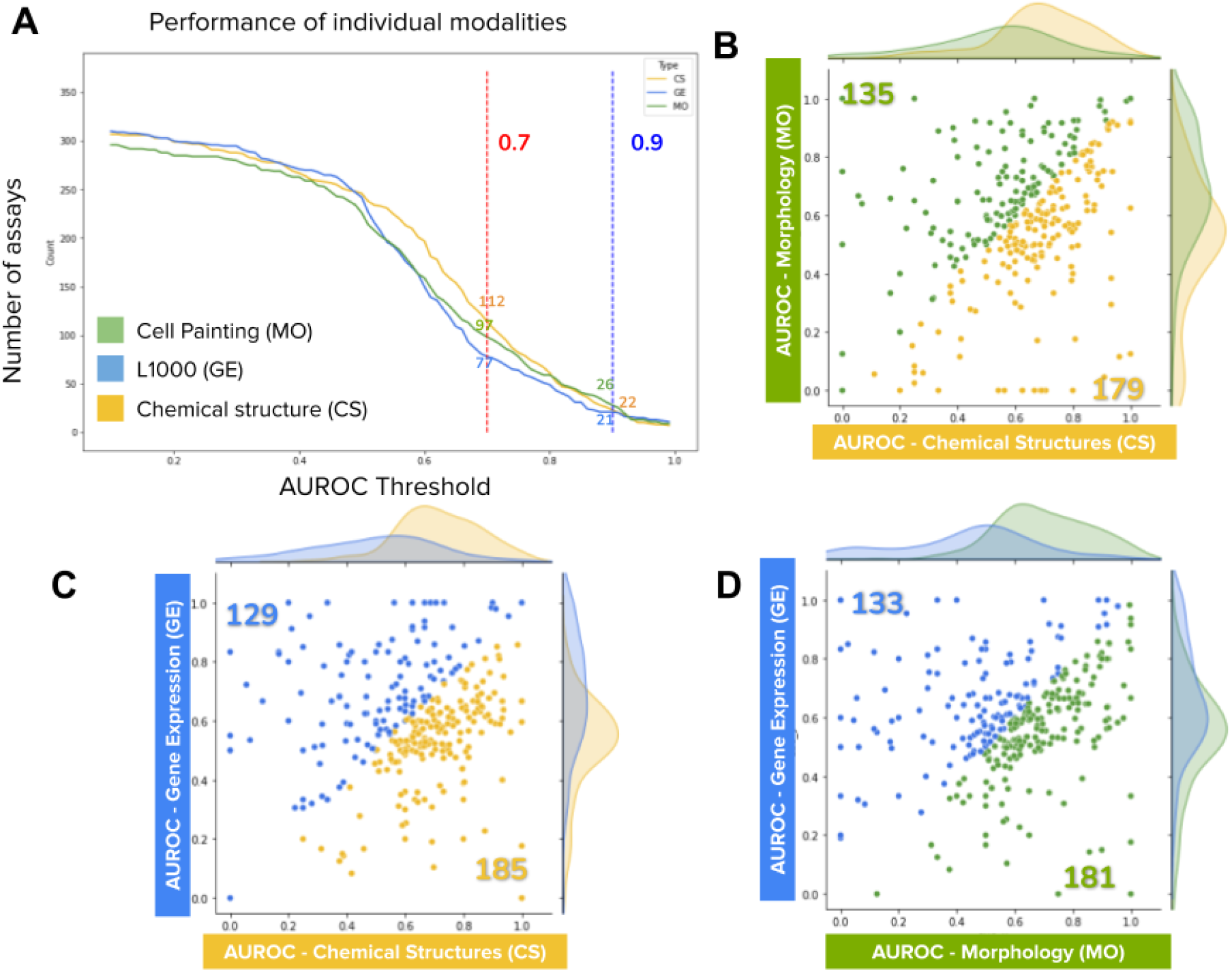
Area under the curve (AUROC) performance of the three individual modalities evaluated in our study: Chemical Structures (CS), Gene Expression (GE), and Morphology (MO). A) Number of assays predicted by each modality at specific AUROC thresholds. As the AUROC threshold is increased, the number of assays meeting the threshold decreases for all modalities. The two thresholds discussed in this paper are highlighted in red (0.7) and blue (0.9). CS outperforms MO and GE until about the 0.8 AUROC threshold, and at higher thresholds MO takes over. B, C, D) Scatter plots of AUROC for pairs of modalities. Each point in the plots represents an assay, the x coordinate indicates the AUROC obtained in one modality, and the y axis represents the AUROC obtained in the other modality. Colors represent the three individual modalities: CS (yellow), GE (blue) and MO (green). Points (assays) above or below the diagonal (equal performance) are colored according to the modality that has the highest AUROC. The two colored numbers inside the plot indicate the total number of assays with higher AUROC with respect to the other modality in the same plot. The counts of points indicate the number of assays where one modality is better than the other. Note that there are many points far off the diagonal, indicating high AUROC in one modality but low in the other. This indicates potential for complementary and fusion among the different data modalities.

**Supplementary Figure 5.**
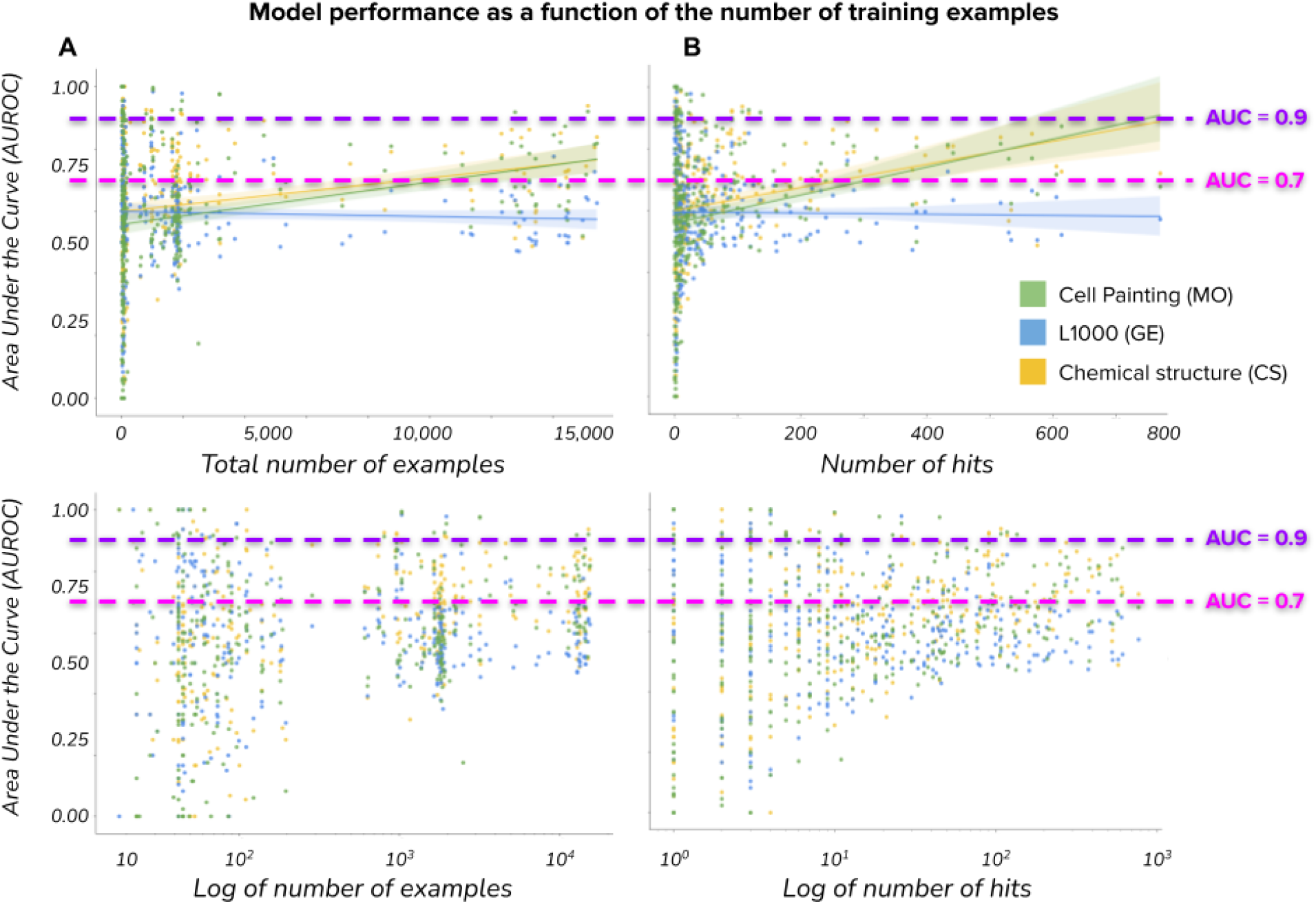
The performance of predictive models is slightly correlated with the number of available training examples; several assays can be predicted with high accuracy (AUROC > 0.9) using only a few example hits (points above the purple line). The plots show on the vertical axis the test set accuracy as a function of (A) the total number of example readouts, and (B) the number of hits available for training. Plots in the bottom row show the same data with log scale in the horizontal axis to highlight the trend with few examples. Each point is an assay predictor and its color indicates what data modality was used for training it. Note that assay prediction accuracy can vary from very low to very high with a small number of training examples, indicating that performance depends on the specific activity measured by the assay.

### Folds of improvement

**Supplementary Figure 6.**
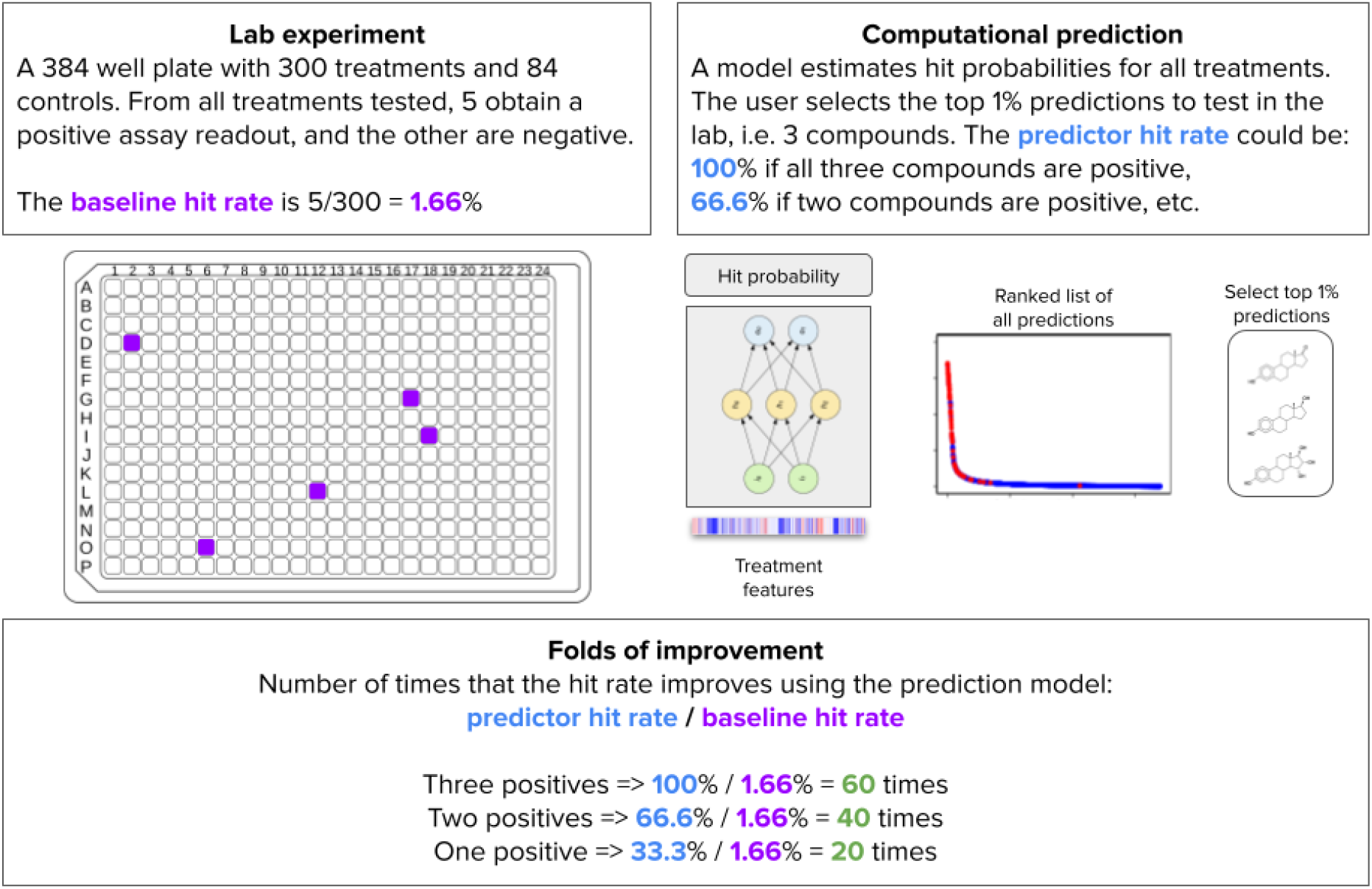
Illustration of the “Folds of improvement” metric. The example assumes a chemist testing a set of 300 candidate compounds where only 5 of them are positive hits. The ratio of hits vs tested compounds is a rough estimate of the probability of finding a hit by chance. A pre-trained computational predictor could rank the same compounds in silico from high probability of being a hit to low probability. We simulate the case where the chemist only selects the top 1% predictions for further wet lab testing, which is a reasonable cut off in real world high-throughput screens with very large compound libraries. By estimating the ratio of hits found in the top 1% subset that is actually tested in vitro, we then compute the folds of improvement as the ratio of the hit rates in each approach. Folds of improvement can be understood as the number of times that the experimental efficiency improves by using a predictor to filter unlikely hits and bring promising candidates to the top of the list.

**Supplementary Figure 7.**
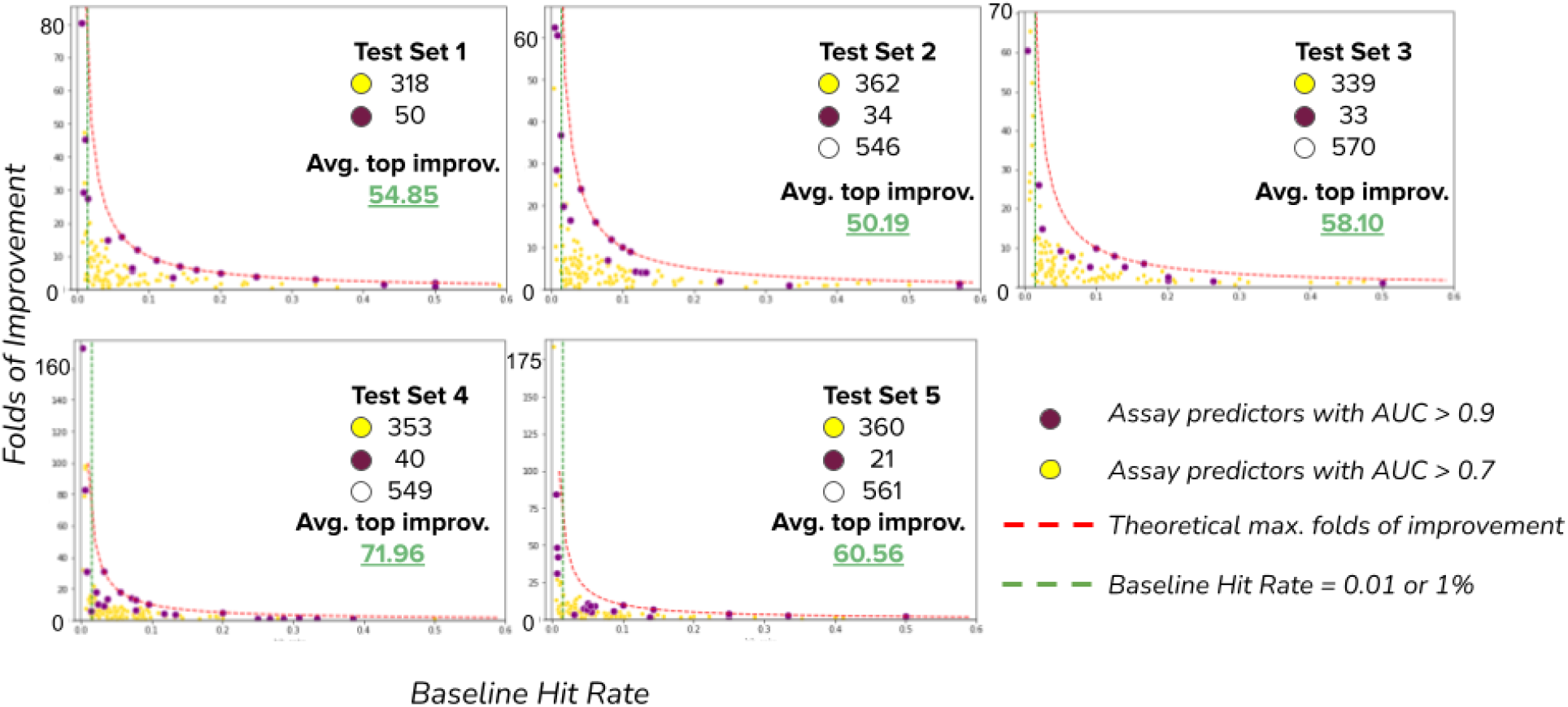
Improvement of hit rates for the assays in the dataset. Each plot corresponds to the results in one split of the 5-fold cross-validation experiment (see Supplementary Figure 1). The points in the plots represent one assay predictor that uses one of the three data modalities (CS, GE or MO) or combinations of them. Assay predictors with AUROC > 0.7 are displayed in yellow and predictors with AUROC > 0.9 are displayed in purple. Assay predictors with AUROC < 0.7 are not shown. The horizontal axis represents the baseline hit rate, i.e., the proportion of compounds found to be hits in the set of tested compounds for an assay (see Supplementary Figure 6). The vertical axis presents the folds of improvement of assay predictions obtained with a machine learning predictor as a function of the baseline hit rate. Accurate predictors (AUROC > 0.9) often offer improvements up to the theoretical maximum (100% divided by the assay’s baseline hit rate), and higher-fold improvements are only possible for assays with a lower baseline hit rate, i.e. with rare hits.

### Data fusion

**Supplementary Figure 8.**
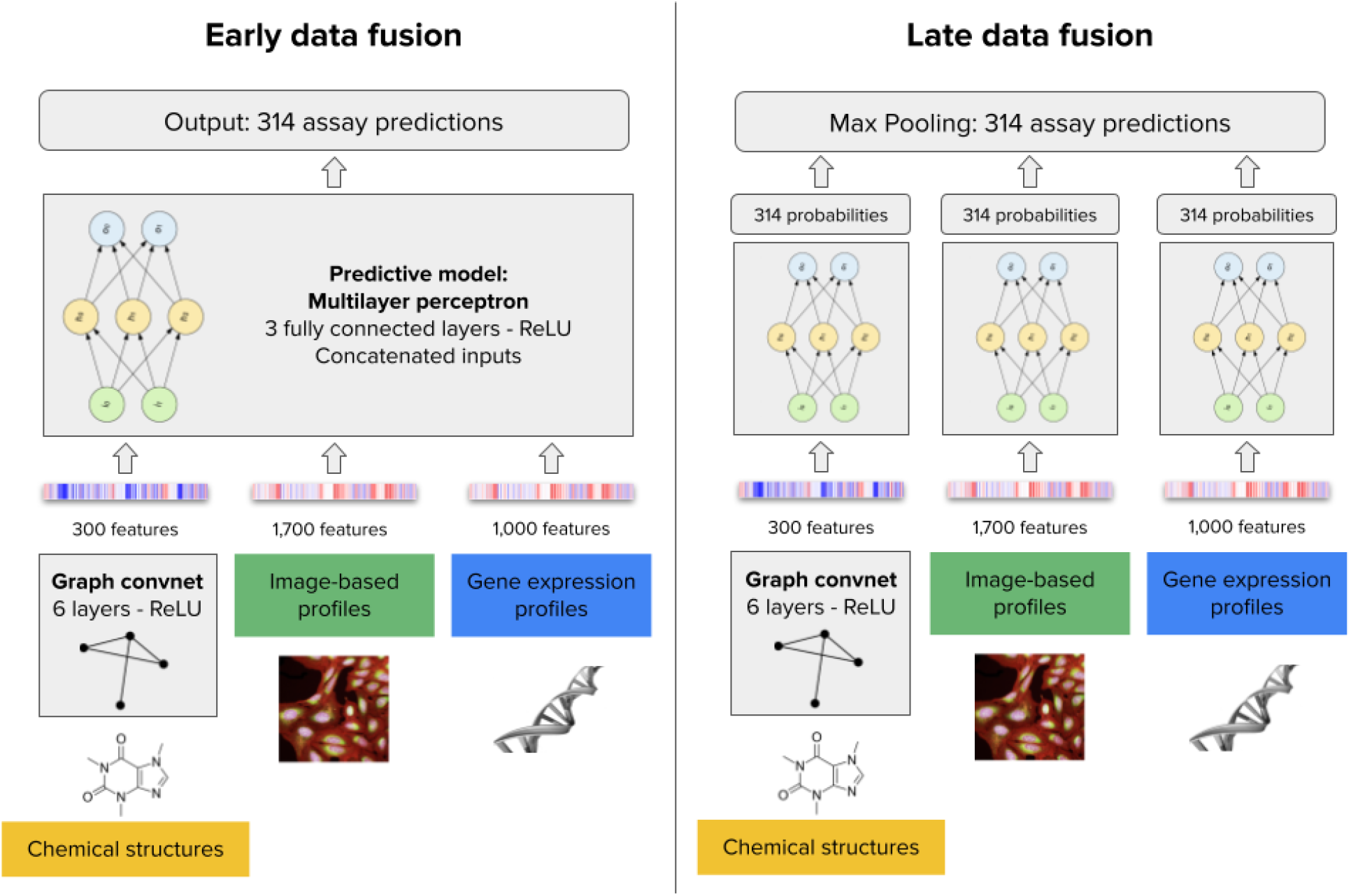
Architecture of early and late data fusion models. The early data fusion model takes the three data modalities as input by obtaining features from each and then concatenating their representations. The architecture is a multilayer perceptron with three fully connected layers, 2,000 input features and 314 output predictions. The late data fusion model has one multilayer perceptron with three fully connected layers independently for each data modality. The three feature vectors are analyzed separately to produce 314 output probabilities in each case, which are later aggregated with a max-pooling operator to reduce them into a single vector of 314 assay predictions.

Combining disparate data modalities is a computational challenge especially when not all the assays can be accurately predicted. Our results indicate that the three data modalities only predict a small fraction of the assays in common (Figure 2B, only three assays are predicted by all modalities), suggesting that in many cases, at least one of the data modalities is effectively introducing noise for predicting a given assay. When one of the data modalities cannot signal the bioactivity of interest, the noise-to-signal ratio in the feature space increases, making it more challenging for predictive models to succeed. This explains why late fusion, which independently looks at each modality, tends to preserve better performance.

We expanded our discussion about data fusion, but we also limited the scope of our current experiments to exploring simple approaches. In future research, we hope that the community or our own team comes up with better methodologies to better model this problem.

**Supplementary Table 1.**
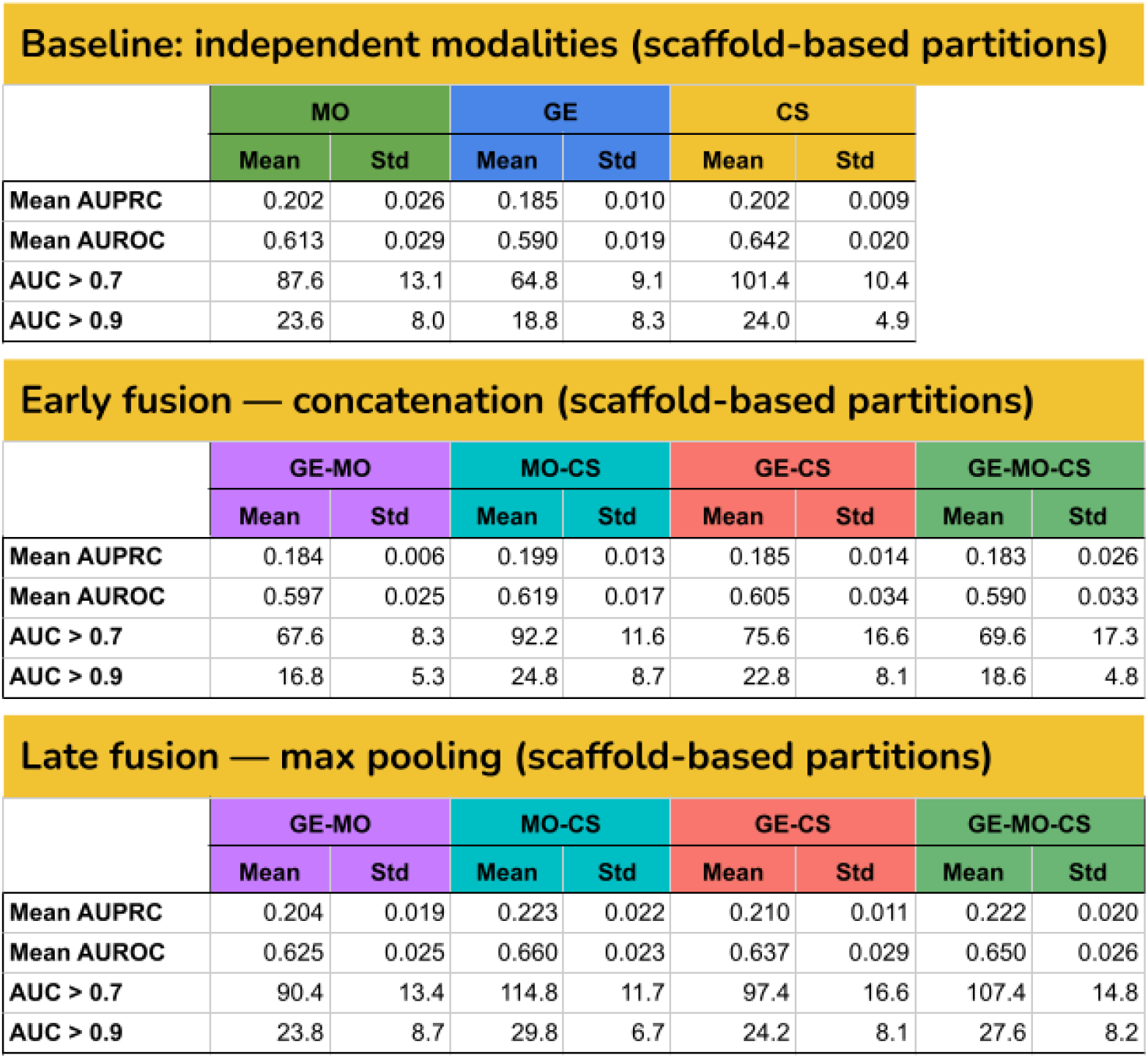
Overall performance of profiling modalities and their combinations presented in the columns of the tables. Early fusion refers to concatenation of feature vectors before training predictive models, while late fusion refers to keeping the maximum prediction of individual models (see Supplementary Figure 8). The tables present four performance metrics in the rows: Mean AUPRC, mean AUROC, number of assays predicted with AUROC > 0.7, and number of assays predicted with AUROC > 0.9. For each experiment, we obtain the mean and standard deviation of the metric. In the case of the mean value for all metrics, higher numbers indicate better performance. Late fusion yields the largest number of predictors with AUROC > 0.9 overall, and also for all combinations of descriptors.

Baseline: independent modalities (scaffold-based partitions)
|  | MO |  | GE |  | CS |  |
| --- | --- | --- | --- | --- | --- | --- |
|  | Mean | Std | Mean | Std | Mean | Std |
| Mean AUPRC | 0.202 | 0.026 | 0.185 | 0.010 | 0.202 | 0.009 |
| Mean AUROC | 0.613 | 0.029 | 0.590 | 0.019 | 0.642 | 0.020 |
| AUC > 0.7 | 87.6 | 13.1 | 64.8 | 9.1 | 101.4 | 10.4 |
| AUC > 0.9 | 23.6 | 8.0 | 18.8 | 8.3 | 24.0 | 4.9 |

Early fusion — concatenation (scaffold-based partitions)
|  | GE-MO |  | MO-CS |  | GE-CS |  | GE-MO-CS |  |
| --- | --- | --- | --- | --- | --- | --- | --- | --- |
|  | Mean | Std | Mean | Std | Mean | Std | Mean | Std |
| Mean AUPRC | 0.184 | 0.006 | 0.199 | 0.013 | 0.185 | 0.014 | 0.183 | 0.026 |
| Mean AUROC | 0.597 | 0.025 | 0.619 | 0.017 | 0.605 | 0.034 | 0.590 | 0.033 |
| AUC > 0.7 | 67.6 | 8.3 | 92.2 | 11.6 | 75.6 | 16.6 | 69.6 | 17.3 |
| AUC > 0.9 | 16.8 | 5.3 | 24.8 | 8.7 | 22.8 | 8.1 | 18.6 | 4.8 |

**Supplementary Table 1.** Overall performance of profiling modalities and their combinations presented in the columns of the tables.
|  | GE-MO |  | MO-CS |  | GE-CS |  | GE-MO-CS |  |
| --- | --- | --- | --- | --- | --- | --- | --- | --- |
|  | Mean | Std | Mean | Std | Mean | Std | Mean | Std |
| Mean AUPRC | 0.204 | 0.019 | 0.223 | 0.022 | 0.210 | 0.011 | 0.222 | 0.020 |
| Mean AUROC | 0.625 | 0.025 | 0.660 | 0.023 | 0.637 | 0.029 | 0.650 | 0.026 |
| AUC > 0.7 | 90.4 | 13.4 | 114.8 | 11.7 | 97.4 | 16.6 | 107.4 | 14.8 |
| AUC > 0.9 | 23.8 | 8.7 | 29.8 | 6.7 | 24.2 | 8.1 | 27.6 | 8.2 |

### Data modalities

**Supplementary Figure 9.**
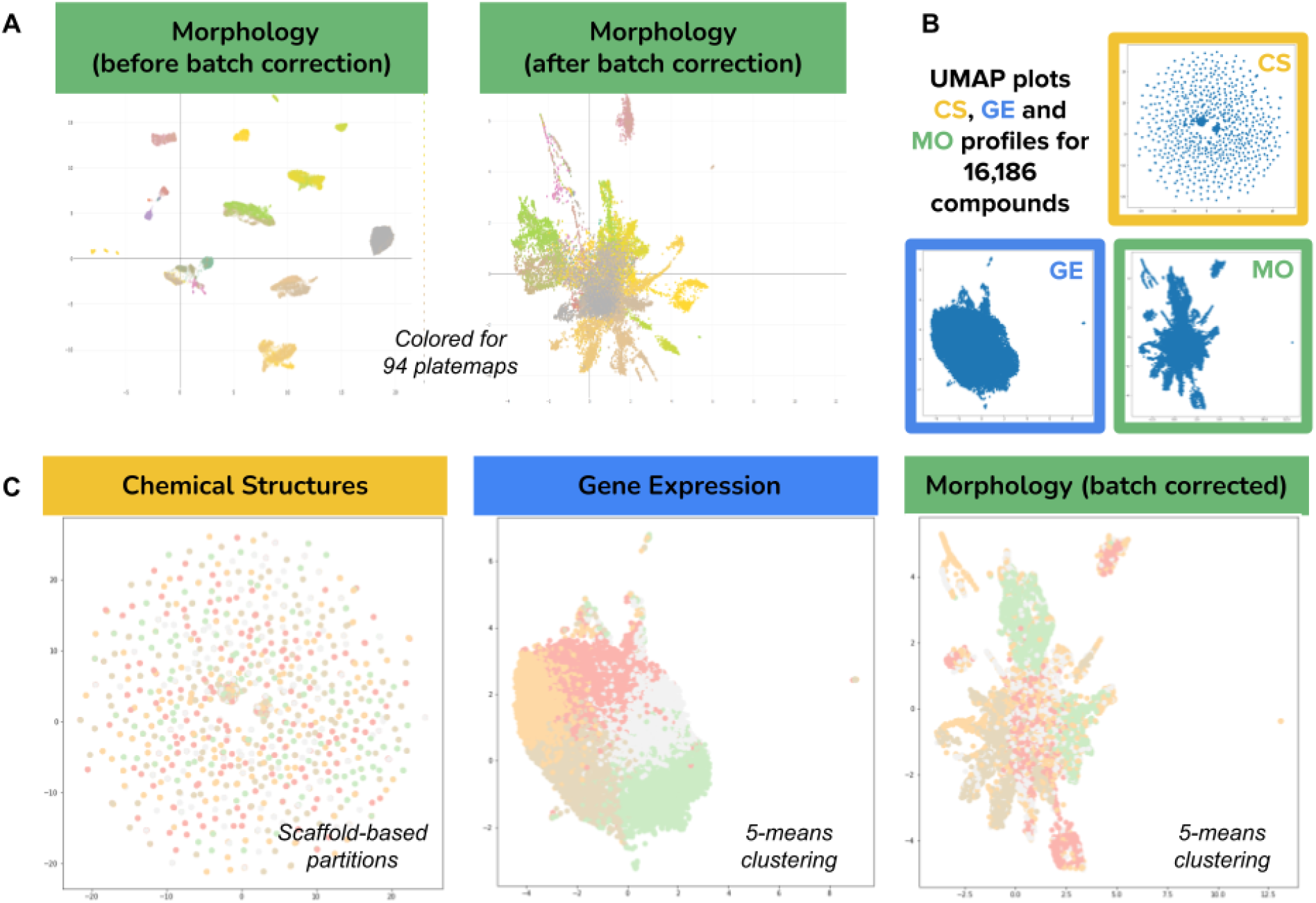
Compound embeddings in three different feature spaces. Visualization of the high-dimensional feature vectors of all compounds using UMAP projections for the three data modalities used in this work. A) The morphology feature space originally was grouped by technical variation (plate maps), which was corrected using the Typical Variation Normalization (TVN) approach (see Methods) to report all experiments in the manuscript. The color palette for the 94 plate maps is continuous and may have similar tones for consecutive plates. B) Overview of the three feature spaces for all the 16,186 compounds included in the evaluation. Note that chemical structures (CS), gene expression (GE), and morphology (MO), all have very distinctive ways of organizing the signatures of compounds. While CS has many diverse small clusters, GE presents a single cloud, and MO has a central cloud with some medium clusters and branches. C) The same visualization as in B, but colored by clusters obtained for cross-validation experiments (see Supplementary Table 2). We partitioned each feature space using clustering to identify 5 groups for training and test splits. CS was split using Bemis-Murcko clustering, which is based on scaffold similarity, while the corresponding UMAP plot projects data points using the features of the full chemical structure (a different metric, which explains why the colors don’t reveal scaffold clusters). GE and MO were split using k-means clustering, with k=5 for cross-validation in simulated control experiments to determine the influence of the data partition in the results (see Supplementary Table 2).

**Supplementary Table 2.** Results of 5-fold cross-validation control experiments. The tables present the mean results of 5-fold cross-validation experiments according to different data partition policies (see Supplementary Figure 9). The scaffold-based splits reflect the real world scenario more closely, while other split policies are useful as control experiments to identify potential artifacts or biases in the data. For each data modality, we used two encoding versions as follows: MO: original features and batch corrected (BC) features. GE: original features and scaled (S) or renormalized features using the L1 norm. CS: graph convolutional (GC) features and Morgan fingerprints (MF). We use as baseline the results of scaffold-based splits, which are reported in the main text and were used to complete all the analysis in this work. Compared to scaffold-based splits, gene expression and random splits yield slightly higher mean AUROC for all other modalities, which confirms that separating training and test compounds randomly makes the prediction problem easier while not being fully informative in a real setting. Morphology splits decrease performance for all modalities, indicating that the k-means splitting by morphology features (see Supplementary Figure 9) disrupts effective learning by bringing together most compounds of certain assays into only one fold. This can be explained partially by the presence of technical artifacts and by real biological signal that could not be entirely separated with the adopted batch correction method. Finally, the difference in performance between graph convolutional representations of chemical structures and Morgan fingerprints is minor across all experiments. Graph convolutions (CS-GC) have slightly better performance in the real world setting, and comparable performance in other splits. We used GC across all the reported experiments in the main manuscript.

Cluster-based 5-fold cross validation
| Scaffold-based splits — Real world setting |  |  |  |  |  |  |
| --- | --- | --- | --- | --- | --- | --- |
|  | MO | MO-BC | GE | GE-S | CS-GC | CS-MF |
| Mean PR-AUC | 0.203 | 0.202 | 0.185 | 0.178 | 0.202 | 0.201 |
| Mean AUROC | 0.640 | 0.613 | 0.590 | 0.578 | 0.642 | 0.637 |
| AUC > 0.7 | 98.8 | 87.6 | 64.8 | 62.2 | 101.4 | 103.0 |
| AUC > 0.9 | 25.8 | 23.6 | 18.8 | 12.6 | 24.0 | 25.2 |

| Gene expression splits (simulation) |  |  |  |  |  |  |
| --- | --- | --- | --- | --- | --- | --- |
|  | MO | MO-BC | GE | GE-S | CS-GC | CS-MF |
| Mean PR-AUC | 0.215 | 0.213 | 0.183 | 0.199 | 0.218 | 0.230 |
| Mean AUROC | 0.655 | 0.637 | 0.590 | 0.586 | 0.659 | 0.668 |
| AUC > 0.7 | 101.0 | 94.4 | 66.4 | 65.6 | 107.8 | 109.4 |
| AUC > 0.9 | 25.8 | 25.6 | 17.2 | 20.2 | 25.0 | 29.8 |

| Morphology(bc)-based splits (simulation) |  |  |  |  |  |  |
| --- | --- | --- | --- | --- | --- | --- |
|  | MO | MO-BC | GE | GE-S | CS-GC | CS-MF |
| Mean PR-AUC | 0.139 | 0.130 | 0.110 | 0.104 | 0.160 | 0.167 |
| Mean AUROC | 0.606 | 0.595 | 0.540 | 0.531 | 0.616 | 0.626 |
| AUC > 0.7 | 58.6 | 47.4 | 34.6 | 32.2 | 67.8 | 71.2 |
| AUC > 0.9 | 11.4 | 9.4 | 8.2 | 5.2 | 13.4 | 17.2 |

**Supplementary Table 2. Results of 5-fold cross-validation control experiments.**
| Random splits (simulation) |  |  |  |  |  |  |
| --- | --- | --- | --- | --- | --- | --- |
|  | MO | MO-BC | GE | GE-S | CS-GC | CS-MF |
| Mean PR-AUC | 0.222 | 0.208 | 0.193 | 0.186 | 0.220 | 0.222 |
| Mean AUROC | 0.654 | 0.623 | 0.599 | 0.582 | 0.662 | 0.663 |
| AUC > 0.7 | 101.6 | 90.2 | 71.8 | 59.2 | 113.2 | 114.0 |
| AUC > 0.9 | 29.0 | 25.2 | 20.4 | 16.6 | 27.6 | 28.6 |

**Supplementary Figure 10.**
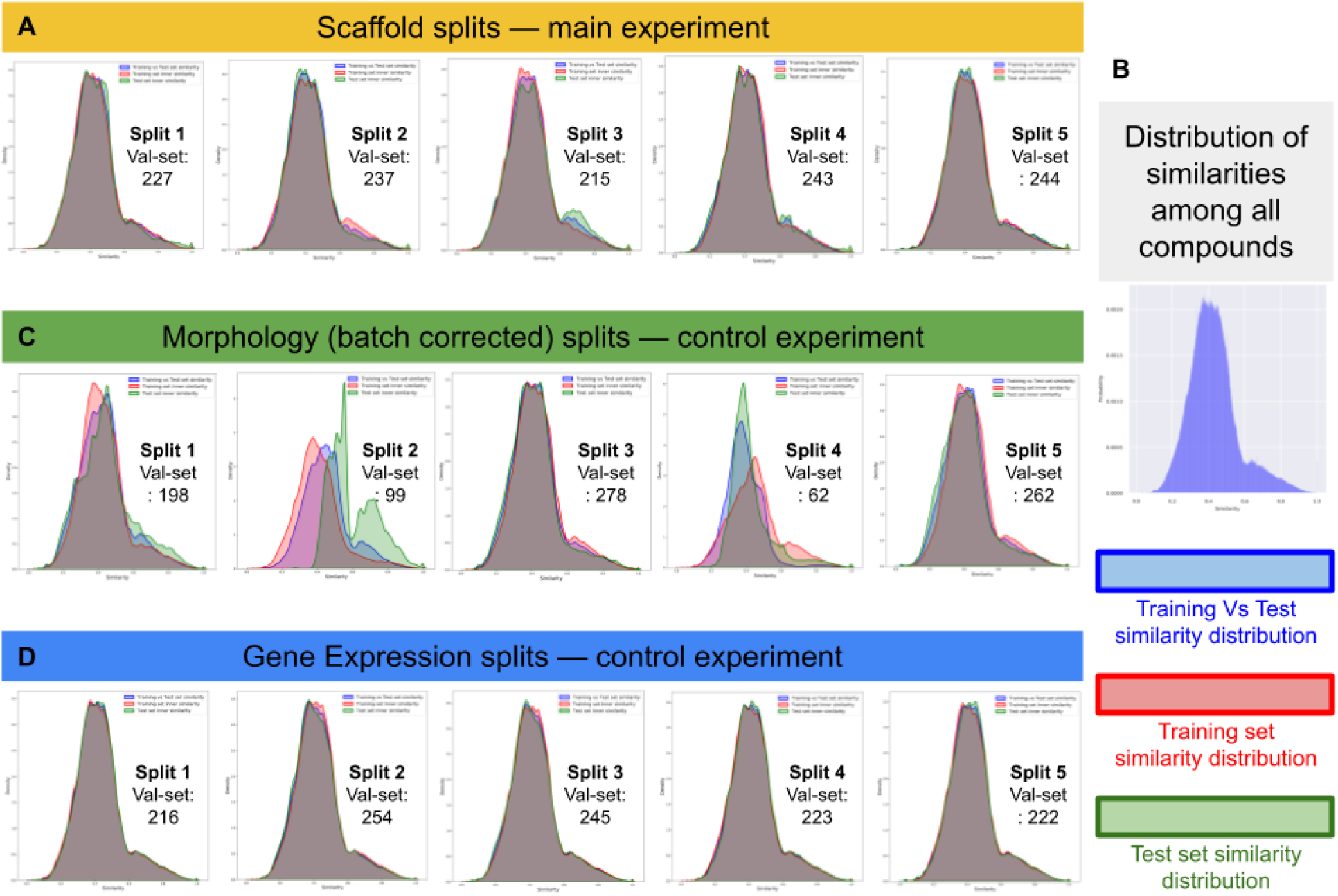
Distribution of compound similarities across training-test splits. We computed the Tanimoto coefficient between Morgan fingerprints of all compounds in the dataset and obtained the distribution of scores (B), which indicates that most compounds are relatively equidistant to each other (consistent with Supplementary Figure 9C). After scaffold-based splitting, this distribution is preserved in training and test partitions in all five folds (A). No major distribution shift is observed with gene-expression splits (D), but two groups in the morphology splits (split 2 and 4) show larger differences likely explained by confounded signal between technical artifacts and biological effects (see Supplementary Table 2 and Supplementary Figure 9).

### Assay data

**Supplementary Figure 11.**
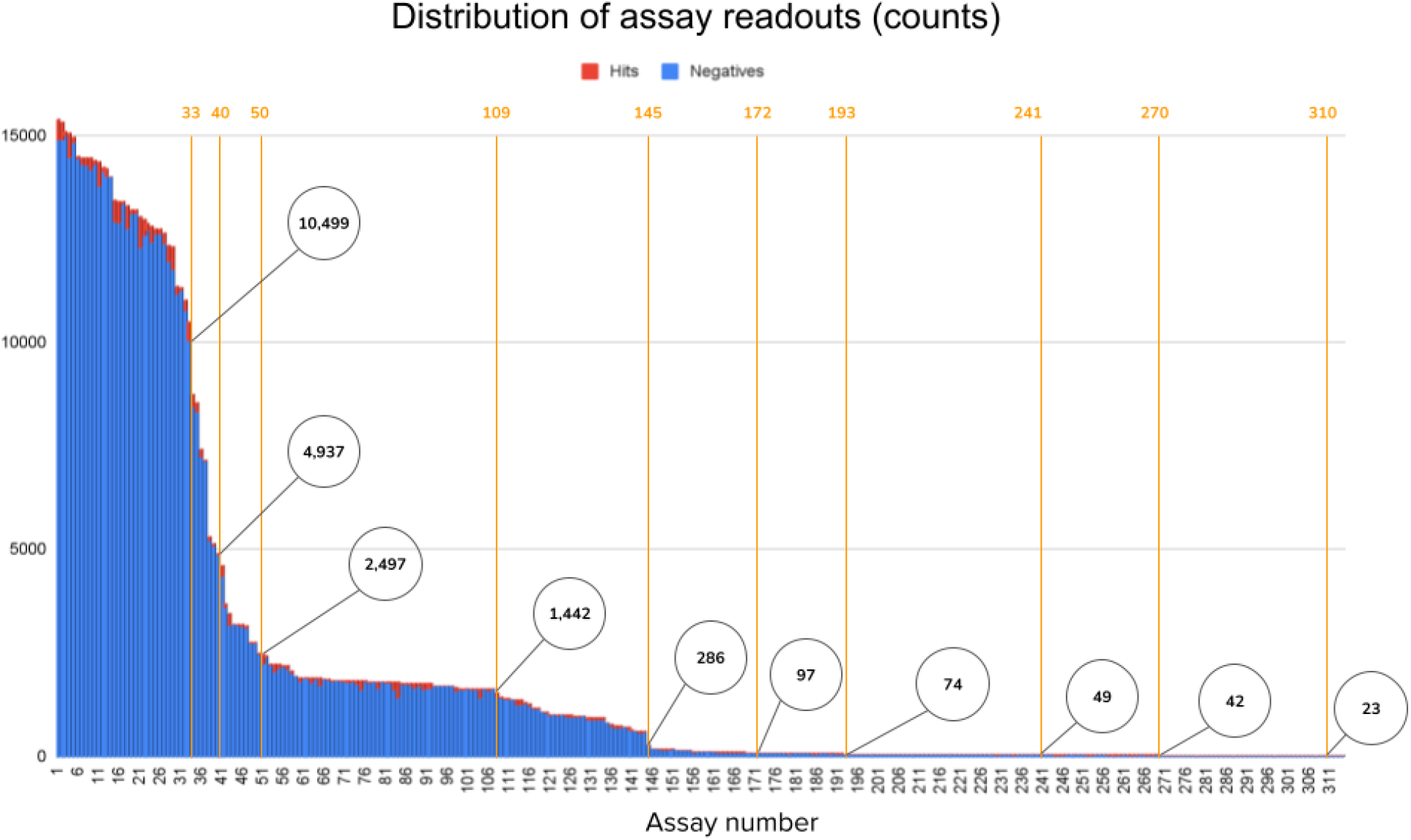
Distribution of assay readouts. The plot shows in the horizontal axis assay identifiers sorted by readout count in decreasing order, and in the vertical axis the count of available readouts for each assay. Readouts can be positive hits (red) or negatives (blue). The circles in the plot indicate the readout count for specific assays in the distribution. Assay readouts follow a long tail distribution, with more than half of the assays having less than a few hundred readouts for training predictive models. Note that the ratio between hits and negative compounds is very small in general (average hit ratio 2.5%).

**Supplementary Figure 12.**
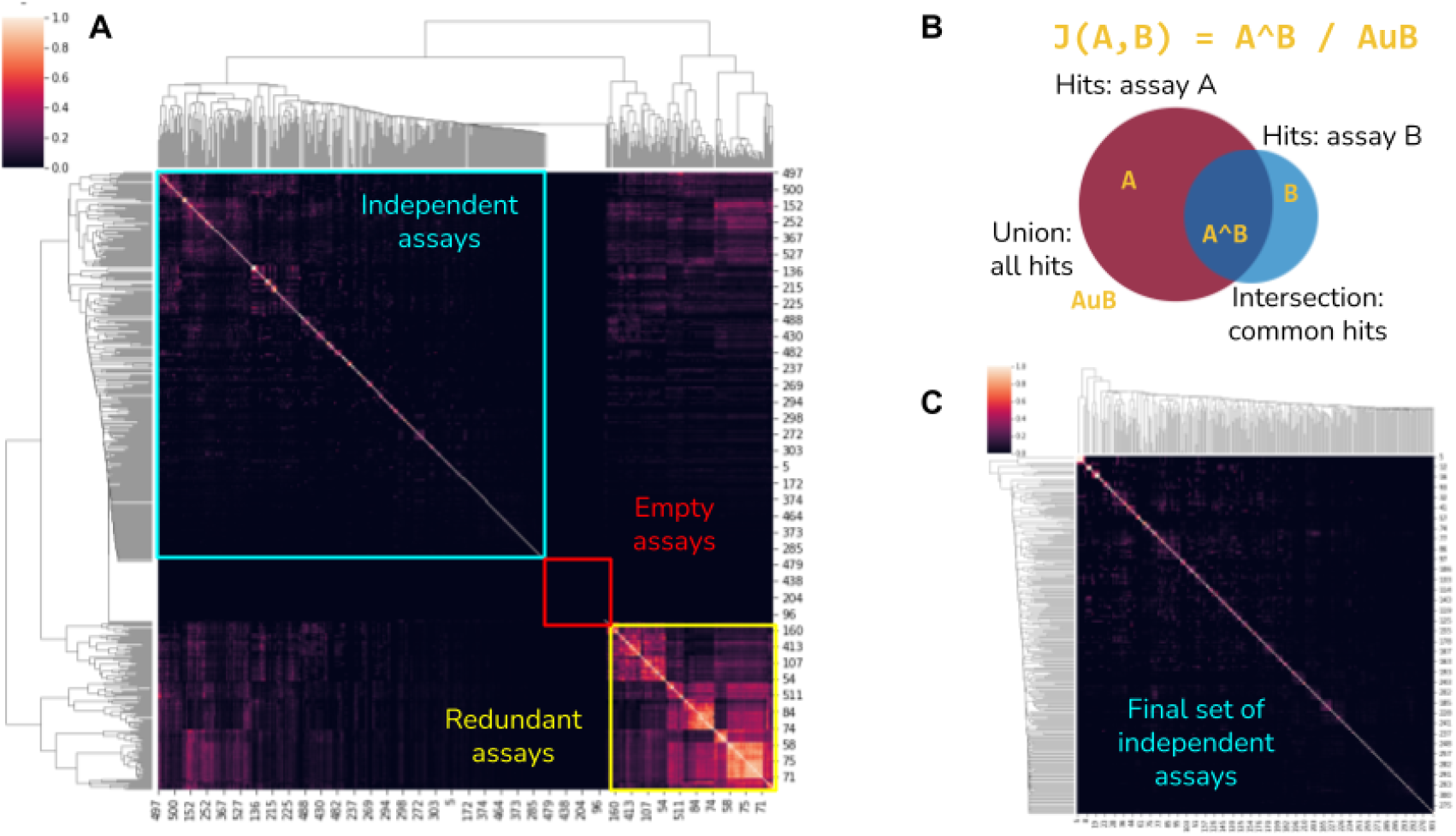
Assay similarity. A) Matrix of assay similarities according to the Jaccard similarity between the sets of positive hit compounds. This matrix presents all the assays originally available for analysis (529). A subset of the assays had no positive hits, and another subset shared a large number of active compounds. We discarded these two groups to preserve a final set of mostly independent assays (shown in panel C). B) Illustration of the Jaccard similarity *J(A,B)* between two assays *A* and *B*. Each assay has a set of positive hits and we compute the ratio of the intersection (hits in common) over the union (count of all total hits) as a metric of similarity between assays. Assays that have many hits in common are likely measuring the same biological activity, and were excluded from our analysis.

**Supplementary Figure 13.**
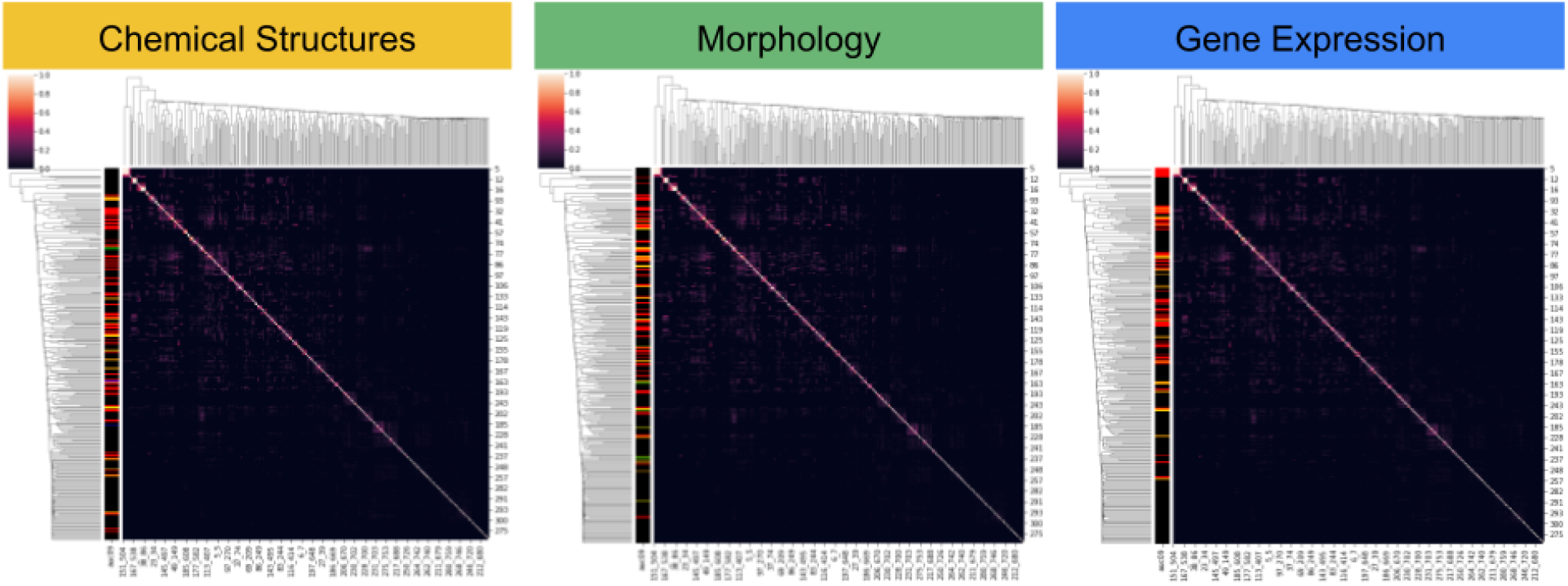
Groups of assays predicted by each modality. The matrix of assay similarities is the same in the three cases: rows and columns are assays and the matrix values are the Jaccard index between the set of hits from two assays. The matrices are clustered in the rows and columns using hierarchical clustering to reveal groups of highly correlated assays. The dendrograms indicate a fairly independent set of assays without major partitions or groupings. The only difference between the matrices is the coloring pattern of the left-hand side bar that indicates whether an assay is correctly predicted by the corresponding modality (chemical structures (CS), morphology (MO), and gene expression (GE)) in any of the cross-validation partitions (non-black colors). This visualization is useful to reveal if the data modalities have preference for making better predictions with certain groups of assays that may have common biological activity. This result indicates that there are no major groups of activation, although accurate predictors tend to be close to each other in the cluster map. The dendrograms reveal a few small assay clusters in the top left of the matrices, and the visualization indicates that only gene expression makes accurate predictions in one of these groups, which may indicate that performance of GE in the experiment may be an overestimate. The other assays are mostly independent from each other according to the Jaccard similarity used in the matrix (see Supplementary Figure 12). We observe that the accuracy patterns in the left of the matrices are mostly randomized, not localized around significant clusters, and they are different from modality to modality.

**Supplementary Figure 14.**
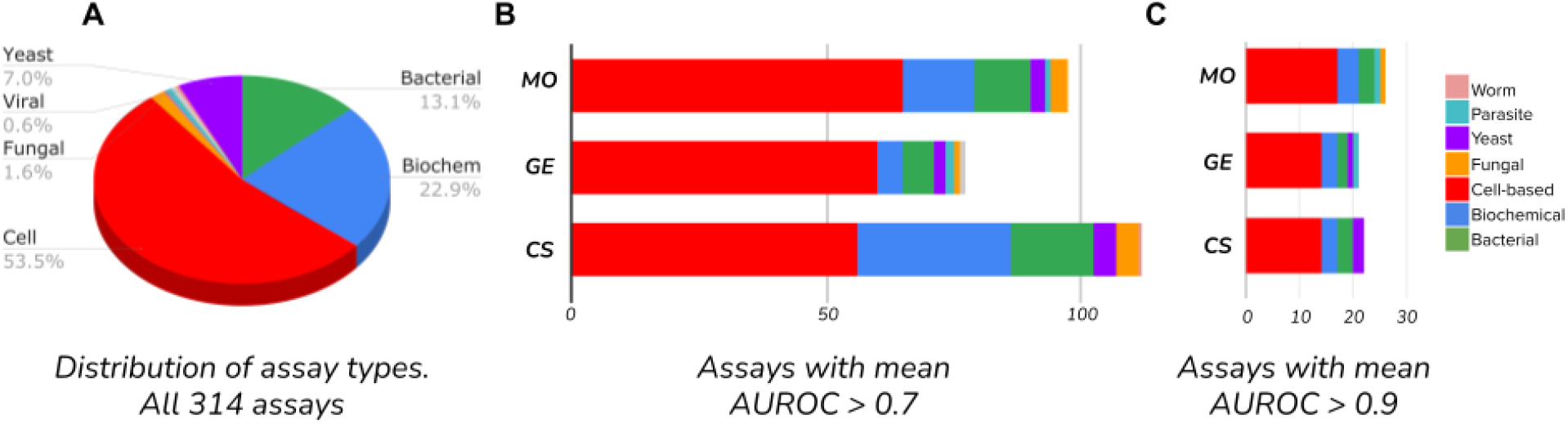
Distribution of assay types as the performance threshold is decreased. The assays used in our study can be one of the seven types listed in the right hand side of the figure. A) Distribution of assays according to their type. B) Distribution of assays that can be predicted with a minimum accuracy of 0.7 AUROC by each of the three data modalities. C) Distribution of assays that can be predicted with a minimum accuracy of 0.9 AUROC by each of the three data modalities. These distributions show that none of the modalities has a strong preference for one type of assay, and that they can predict a diverse array of biological activity.

**Supplementary Table 3.**

| **A** | **0.9 median AUROC** | **CS** | **GE** | **MO** | **CS+GE** | **CS+MO** | **GE+MO** | **CS+GE+MO** | **Evaluated assays** |
| --- | --- | --- | --- | --- | --- | --- | --- | --- | --- |
|  | Cell-based | 8.33% | 8.33% | 10.12% | 10.12% | 10.71% | 8.33% | 10.12% | 168 |
|  | Biochemical | 4.17% | 4.17% | 5.56% | 2.78% | 4.17% | 2.78% | 2.78% | 72 |
|  | Bacterial | 7.32% | 4.88% | 7.32% | 2.44% | 7.32% | 7.32% | 7.32% | 41 |
|  | Yeast | 9.09% | 4.55% | 0.00% | 4.55% | 4.55% | 0.00% | 0.00% | 22 |
|  | Parasite | 0.00% | 50.00% | 50.00% | 0.00% | 50.00% | 50.00% | 50.00% | 2 |
|  | Fungal | 0.00% | 0.00% | 20.00% | 0.00% | 20.00% | 0.00% | 0.00% | 5 |
|  | Viral | 0.00% | 0.00% | 0.00% | 0.00% | 0.00% | 0.00% | 0.00% | 2 |
|  | Worm | 0.00% | 0.00% | 0.00% | 0.00% | 0.00% | 100.00% | 100.00% | 1 |
|  | Homogeneous | 0.00% | 0.00% | 0.00% | 0.00% | 0.00% | 0.00% | 0.00% | 1 |

Supplementary Table 3.
| **B** | **0.7 median AUROC** | **CS** | **GE** | **MO** | **CS+GE** | **CS+MO** | **GE+MO** | **CS+GE+MO** | **Evaluated assays** |
| --- | --- | --- | --- | --- | --- | --- | --- | --- | --- |
|  | Cell-based | 33.33% | 35.71% | 38.69% | 47.02% | 43.45% | 44.64% | 44.05% | 168 |
|  | Biochemical | 41.67% | 6.94% | 19.44% | 26.39% | 40.28% | 16.67% | 29.17% | 72 |
|  | Bacterial | 39.02% | 14.63% | 26.83% | 21.95% | 39.02% | 24.39% | 39.02% | 41 |
|  | Yeast | 22.73% | 9.09% | 13.64% | 31.82% | 22.73% | 13.64% | 27.27% | 22 |
|  | Parasite | 0.00% | 100.00% | 50.00% | 0.00% | 50.00% | 100.00% | 50.00% | 2 |
|  | Fungal | 80.00% | 20.00% | 60.00% | 40.00% | 80.00% | 60.00% | 80.00% | 5 |
|  | Viral | 50.00% | 0.00% | 0.00% | 0.00% | 50.00% | 0.00% | 0.00% | 2 |
|  | Worm | 0.00% | 100.00% | 0.00% | 0.00% | 0.00% | 100.00% | 100.00% | 1 |
|  | Homogeneous | 0.00% | 0.00% | 0.00% | 0.00% | 0.00% | 0.00% | 0.00% | 1 |

